# Humans as Predators of the Biosphere: Technological Modulation of Consumer–Resource Dynamics and Its Implications for Sustainability

**DOI:** 10.64898/2026.04.08.717266

**Authors:** Vanessa P. Weinberger, Nicolás Zalaquett, Mauricio Lima

## Abstract

Humans are just another species on Earth, but modern telecoupled societies and their socioeconomies impose immense consumption demands on the biosphere, detaching from common ecological rules. Starting from a simple ecological consumer–resource model, with humans as the consumers and terrestrial organic carbon (i.e., the biosphere) as the resource, we assume that technology modulates both human carrying capacity, *ν*_0_, and the rate of biosphere consumption, *α*_0_. Three different functional-relation scenarios were tested, modulated by parameter *a*. In all three scenarios, equilibria and stability directly depended on the relative role that technology played in the model parameters, or the *compound technological impact* (*ϵ* ≡ *α*_0_*ν*_0_). Moreover, two of the three scenarios showed Hopf bifurcations and regions with no equilibrium. The models were parameterized and fitted to actual data using a trajectory of more than 150 years. These analyses suggest that we are currently in a stable oscillatory spiral with no immediate Hopf bifurcation threat, but within a trajectory that continuously depletes the biosphere and approaches a collapse in human population size if no changes are made in the relationship that technology has with growth (i.e., *ν*_0_) versus consumption (i.e., *α*_0_) dynamics. Because our predatory dynamics also appear to have shifted from regular predator– prey dynamics toward a supply–demand scenario, with persistently increasing values, the threat of a Hopf bifurcation is now present in our trajectory: changes in the stability of the coexistence equilibrium may arise. This simple model warns that we must pay closer attention to the predatory relations that our technologies are creating with biosphere dynamics, in a way that goes beyond population numbers and technological development alone.

## 1 Introduction

Humans are just another species on Earth and, as such, depend on energy and materials to sustain their metabolism and activities [Brown et al., 2004, Sibly et al., 2012]. However, modern telecoupled societies and their socioeconomies impose immense consumption demands, detaching from common ecological rules and imposing strong pressure on ecological dynamics [Moses and Brown, 2003, Burnside et al., 2012, Brown et al., 2011, Burger et al., 2012]. There is an urgent need to understand the drivers behind increased human population size and consumption, and their implications for sustainable trajectories.

It has been established that humans, through *sociocultural niche construction*, rapidly accumulate cultural traits over time, and their use is capable of altering regular biospheric dynamics at global scales [Ellis, 2015, Steffen et al., 2018, Ellis et al., 2020]. Under the particular sociocultural trajectory that has been selected, human societies have produced the *Technosphere* at the expense of altering ecosystem dynamics at all spatiotemporal scales [Søgaard Jørgensen et al., 2023].

*Technology* is cultural information “capable of enhancing the behavioral capacity of organisms beyond their biological capabilities” [Ellis, 2015], and it evolves rapidly because it changes within and across generations and accumulates through time through the *ratchet effect* [Arthur, 2009, Ellis, 2015]. This can create a *runaway evolutionary* dynamic: technological adoption requires new cultural adaptations that then require further *innovations* or cultural evolution, generating a positive feedback loop of increasing enhancement and dependence [Bettencourt et al., 2007, Ellis, 2015, Weinberger et al., 2017, Lima et al., 2023, Richerson et al., 2023]. However, this runaway and accelerated dynamic has been argued to lie at the core of human unsustainability, because it outpaces regular systemic adaptation [Snyder, 2020, Lenton et al., 2021, Weinberger et al., 2023].

A clear expression of this unsustainable technological process is the recurrent expansion of energy access in modern human societies, together with increases in the scale and efficiency of energy throughput achieved through societal development [Bettencourt et al., 2007, Ellis, 2015, Ellis et al., 2020, Snyder, 2020, Lenton and Scheffer, 2023, Snyder, 2023, Lima, 2025]. From fire use and domestication to industrial reliance on *extra-metabolic* energy (coal, gas, and oil), these technological *innovations* expanded human capacity to appropriate biological energy and perform work [Haberl et al., 2007, Haberl et al., 2014, Brown et al., 2011, Schramski et al., 2015, Kleidon, 2024]. In doing so, humans not only increased in population size, but also increased their per capita energy demand through expanding socioeconomies, or *social metabolism* [Brown et al., 2011, DeLong et al., 2010, Burger et al., 2012]. It is argued that population, economics and energy consumption growths cannot be dissociated from ecosystem degradation, such as climate change, biodiversity loss, soil erosion, and ocean acidification [Barnosky et al., 2012, Steffen et al., 2018]. In fact, it represents the explicit manifestation of the interrelated interrelations between humans and the natural systems that cannot be resolved within the dominant narrative of environmentally sustainable growth [Odum and Odum, 2006, Herrmann-Pillath, 2015, Herrmann-Pillath, 2016, Herrmann-Pillath, 2025, Vermeij, 2023].

Given this background, in this research we modify a simple ecological consumer–resource model to couple key biospheric processes with human population growth and societal development, and evaluate sustainability under technology-modulated scenarios. Recognizing that technology can expand human carrying capacity while modulating the per capita rate of terrestrial organic carbon consumption (proxy for biosphere consumption), we determine the technology-modulated parameter space in which biologically relevant stable coexistence equilibria can exist. Section 2 describes the model. Section 3 presents the analytical and fitting results, and section 4 discusses them. Finally, section 5 concludes.

## 2 The Model

All biological work is supported by biochemical reactions that continually transform molecules and obtain energy in the process (i.e., metabolisms). This set of organic energy reactions is what we define as the biosphere: all biological energy transformation processes that occur on planet Earth. Humans, as part of the biosphere, are intrinsically embedded within this intricate network of energy relations, and any increase in their so- ciocultural energy demands necessarily implies support from the biosphere. This is the framework on which we base our model.

We model the human–biosphere interaction through an ecological consumer–resource dynamic, with *X* the number of consumer individuals (humans) and *Y* the resource (i.e. the biosphere). We used the terrestrial organic carbon production –a major organic molecule for metabolisms and biomass and for which multiple quantifications exist– as proxy of the biosphere. We are aware that is only one component of the biogeochemical carbon cycle, and only one component of all biospheric processes. However, terrestrial organic carbon is responsible for at least 50% of organic carbon production and retention, and have been greatly impacted by agricultural, livestock and mining activities.

Technology (*Z*) has two main roles within this human–biosphere interaction: it affects (i) human carrying capacity, adjusting the number of human individuals that can be supported per unit of organic carbon (parameter *ν*_0_(*Z*^∗^)); and (ii) per capita organic carbon depletion rates (parameter *α*_0_(*Z*^∗^)). Social metabolism is indirectly included in these parameters. It should affect the relation that technology has with (i) *ν*_0_, where increases reflect investment in population growth, but decreases are interpreted as greater per capita sociocultural energy costs for maintaining a human individual in that society; and (*α*_0_), where sustainable outcome only relate to efficiency increases in this relation, as any other increase implies increasing pressure on biosphere stocks. However, modern technological modes of biosphere appropriation do not seem to decouple from increasing organic energy consumption.

Notice that we do not model technology as a dynamic state variable, but as a fixed technological stock (*Z*^∗^) that affects model parameters and is assumed to change across historical periods.

The proposed general human–biosphere model is

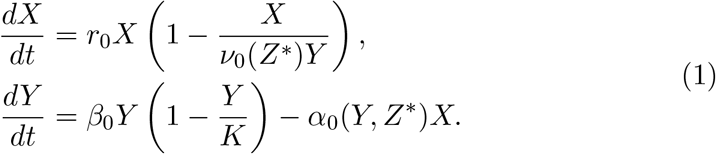

Here, the human population growth equation follows logistic dynamics, with *r*_0_ the maximum biological annual population growth rate, and *ν*_0_ a parameter representing how many human individuals can be allocated per unit of organic carbon for a given fixed technological stock *Z*^∗^.

We also assume that the stock of organic energy grows logistically, with *β*_0_ the turnover rate of organic carbon and *K* its carrying capacity. Humans access organic carbon stocks through different ecological consumer–resource functional relations, depleting them at per capita rates *α*_0_(*Y, Z*^∗^). Different functional relations are used, representing alternative consumer–resource scenarios, modulated by parameter *a*:

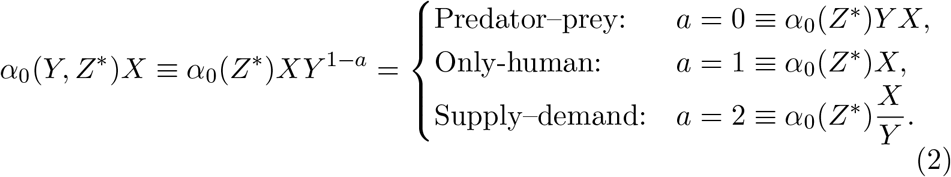

For the predator–prey scenario (*a* = 0), we assume that humans interact with the biosphere through a mean-field encounter relation, eliminating any explicit spatiotemporal component in rates of encounter and energy processing. For the only-human scenario (*a* = 1), we assume that the functional relation does not depend on organic carbon stock, but only on human processing capacity. Finally, for the supply–demand scenario (*a* = 2), we assume that humans exploit the biosphere relative to its availability. These scenarios allow us to evaluate contrasting exploitation dynamics and their sustainability implications.

## 3 Results

### 3.1 Analytical results

Details of the analytical resolution are contained in the supplementary material. Here we present the key analytical results for each functional-relation scenario. For readability, we drop the dependence on technological stock *Z*^∗^ in the equations below.

Irrespective of the functional relation used, the two trivial equilibria are the same for all scenarios. The first, **human extinction** (*X*^∗^ = 0, *Y* ^∗^ = *K*), is an unstable saddle point with one positive and one negative eigenvalue. The second, **system extinction** (*X*^∗^ = 0, *Y* ^∗^ = 0), requires special care, because its stability behavior depends on the trajectory of approach due to indeterminacies and singularities [Kot, 2001, Safuan et al., 2012].

Nontrivial equilibria and their stability change with each functionalrelation scenario.

#### 3.1.1 Case *a* = 0: Predator–prey functional relation

The dimensional model is

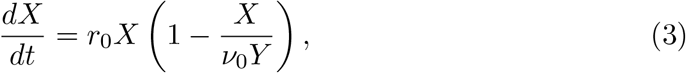

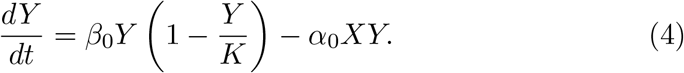

Using the substitutions

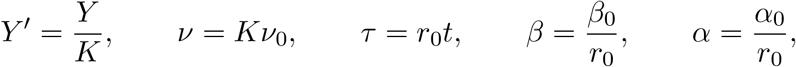

the system becomes

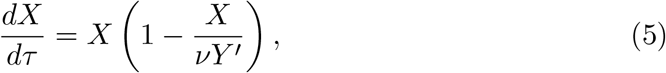

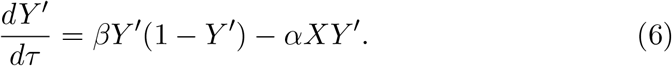

This system has one nontrivial equilibrium:

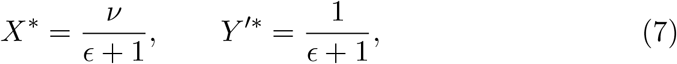

with *ϵ* = *αν/β*.

Analysis of the Jacobian determinant and trace at this equilibrium shows that it is always stable for positive parameter values, with no Hopf bifurcation and no limit cycles. However, the discriminant shows a transition between a stable node and a stable spiral, determined by the critical curve

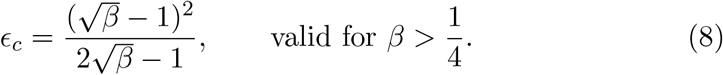

Figure 1A shows the stability diagram for this equilibrium in the (*β, ϵ*) parameter space.

**Figure 1:**
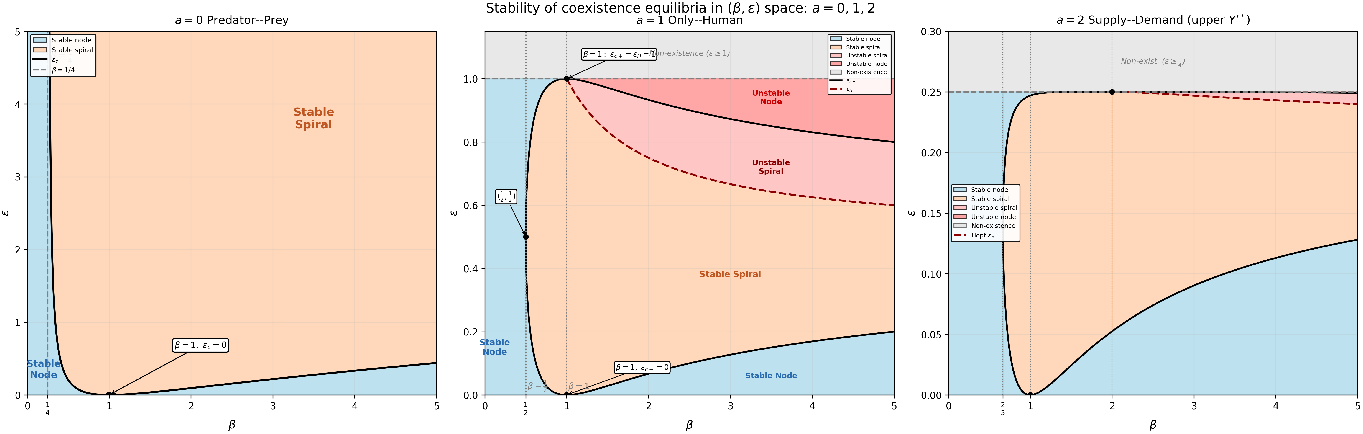
Stability diagram for the nontrivial fixed point under all functional-relation scenarios.

In general, whenever 0 *< β <* 1*/*4, the equilibrium is a stable node. If *β >* 1*/*4, the stable node persists whenever 0 *< ϵ < ϵ*_*c*_; otherwise, the equilibrium becomes a stable spiral.

#### 3.1.2 Case *a* = 1: Only-human functional relation

The dimensional model is

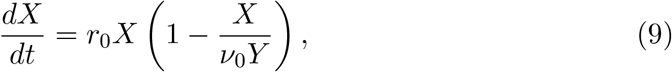

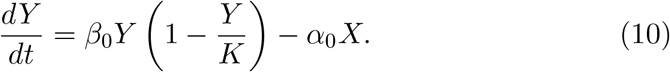

Using the substitutions

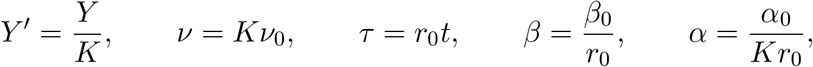

the system becomes

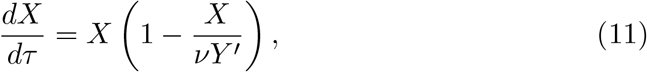

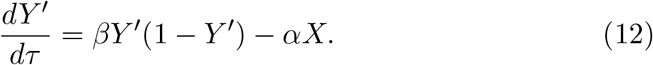

This model also contains a single nontrivial equilibrium:

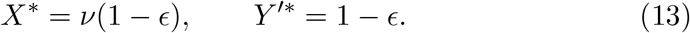

For biological feasibility, 0 *< ϵ <* 1 is required.

The determinant of the Jacobian at this equilibrium is always positive for *ϵ* ∈ (0, 1), so the equilibrium is never a saddle within its existence range. However, trace analysis shows a Hopf bifurcation threshold at

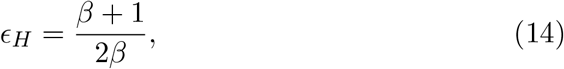

which lies within the valid range only when *β >* 1.

Discriminant analysis shows that the equilibrium behaves as a spiral whenever *ϵ*_*c*−_ *< ϵ < ϵ*_*c*+_, and as a node otherwise, with

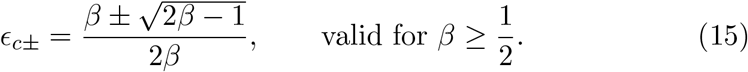

#### 3.1.3 Case *a* = 2: Supply–demand functional relation

The dimensional model is

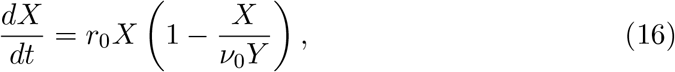

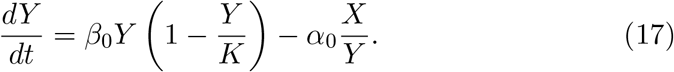

Using the substitutions

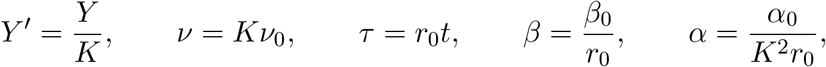

the system becomes

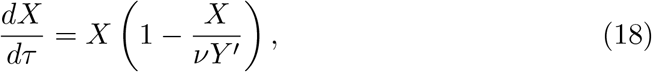

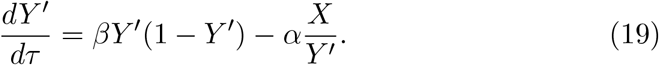

The nontrivial equilibrium is

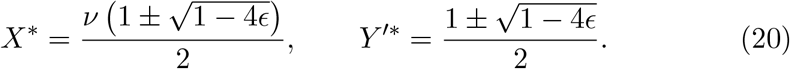

This can generate multiple equilibria depending on *ϵ*. Whenever *ϵ <* 1*/*4, two distinct equilibria exist; when *ϵ* = 1*/*4, a single equilibrium appears; and when *ϵ >* 1*/*4, no real equilibrium exists.

The lower equilibrium is always a saddle point. The upper equilibrium is never a saddle point, and trace analysis shows a Hopf bifurcation at

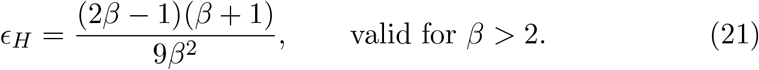

A spiral region exists for *β >* 2*/*3, vanishing at

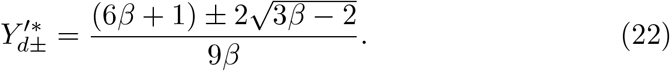

### 3.2 Parameters and model fitting

Dimensional parameters needed for the model were obtained from the scientific literature. Details are provided in the supplementary material. The adopted values place the system in the dynamically rich regime of the stability diagrams, with

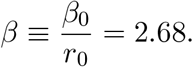

Global human population from 1850 to 2023 and reconstructed terrestrial global carbon stock were used for fitting. Because technology-modulated parameters *α*_0_ and *ν*_0_ are expected to change across the *>* 150-year record due to technological and socioeconomic shifts, we modeled the time series as piecewise-constant functions over *N*_bp_ + 1 segments separated by *N*_bp_ breakpoints.

The predator–prey functional relation selected four breakpoints (1911, 1954, 1969, 1993), corresponding approximately to the Industrial Revolution, the Great Acceleration, the Green Revolution, and the Digital Revolution. The other functional-relation scenarios repeated these dates and additionally selected years around the First and Second World Wars (see Table 3).

**Table 1:** Model variables and parameters.

| Symbol | Description | Units |
| --- | --- | --- |
| *State variables* |  |  |
| $X$ | Human population | ind |
| $Y$ | Biosphere stock (organic carbon) | C |
| $Z^*$ | Technological stock (fixed) | – |
| $t$ | Time | T |
| *Dimensional parameters* |  |  |
| $r_0$ | Intrinsic human population growth rate | $T^{-1}$ |
| $\beta_0$ | Biosphere turnover rate | $T^{-1}$ |
| $K$ | Biosphere carrying capacity | C |
| $\nu_0$ | Human allocation per unit of organic carbon | $\text{ind C}^{-1}$ |
| $\alpha_0$ ($a = 0$) | Per-capita consumption rate (predator–prey) | $\text{ind}^{-1} T^{-1}$ |
| $\alpha_0$ ($a = 1$) | Per-capita consumption rate (only-human) | $\text{C ind}^{-1} T^{-1}$ |
| $\alpha_0$ ($a = 2$) | Per-capita consumption rate (supply–demand) | $\text{C}^2 \text{ind}^{-1} T^{-1}$ |
| *Dimensionless quantities* |  |  |
| $Y' = Y/K$ | Normalized biosphere state | – |
| $\tau = r_0 t$ | Dimensionless time | – |
| $\beta = \beta_0/r_0$ | Relative biosphere-to-human growth rate | – |
| $\epsilon = \alpha\nu/\beta$ | Compound technological impact | – |

**Table 2:** Parameter values adopted for the fitted model and their basis.

| Parameter | Value used | Reference / basis |
| --- | --- | --- |
| $K$ | 916 GtC | Potential vegetation under current climate without humans |
| $\beta_0$ | $0.067 \text{ yr}^{-1}$ | Inverse of carbon turnover rate at equilibrium |
| $r_0$ | $0.025 \text{ yr}^{-1}$ | Conservative value for hunter-gatherers |
| $\beta$ | 2.68 | $\beta = \beta_0/r_0$ |

**Table 3:**
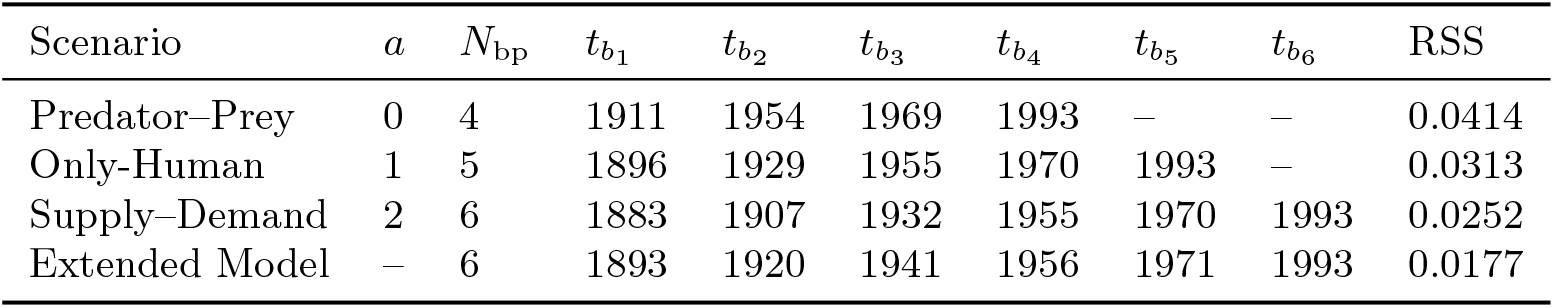
Breakpoint years selected for each individual model scenario.

| Scenario | $a$ | $N_{bp}$ | $t_{b_1}$ | $t_{b_2}$ | $t_{b_3}$ | $t_{b_4}$ | $t_{b_5}$ | $t_{b_6}$ | RSS |
| --- | --- | --- | --- | --- | --- | --- | --- | --- | --- |
| Predator–Prey | 0 | 4 | 1911 | 1954 | 1969 | 1993 | – | – | 0.0414 |
| Only-Human | 1 | 5 | 1896 | 1929 | 1955 | 1970 | 1993 | – | 0.0313 |
| Supply–Demand | 2 | 6 | 1883 | 1907 | 1932 | 1955 | 1970 | 1993 | 0.0252 |
| Extended Model | – | 6 | 1893 | 1920 | 1941 | 1956 | 1971 | 1993 | 0.0177 |

**Table 4:** Parameter estimates for the extended model by segment.

| Segment | Time interval | $\nu_0$ | $\alpha_0$ | $\alpha_1$ | $\alpha_2$ |
| --- | --- | --- | --- | --- | --- |
| 1 | 1850–1893 | 0.00296 | $2.170 \times 10^{-2}$ | $5.123 \times 10^{-3}$ | $7.383 \times 10^{-2}$ |
| 2 | 1893–1920 | 0.00469 | $1.971 \times 10^{-2}$ | $6.096 \times 10^{-3}$ | $5.175 \times 10^{-1}$ |
| 3 | 1920–1941 | 0.00683 | $1.728 \times 10^{-2}$ | $5.574 \times 10^{-2}$ | $7.375 \times 10^0$ |
| 4 | 1941–1956 | 0.00931 | $1.224 \times 10^{-2}$ | $9.556 \times 10^{-1}$ | $1.984 \times 10^2$ |
| 5 | 1956–1971 | 0.03317 | $1.168 \times 10^{-2}$ | $1.716 \times 10^{-1}$ | $3.767 \times 10^1$ |
| 6 | 1971–1993 | 0.03802 | $8.035 \times 10^{-3}$ | $4.051 \times 10^{-2}$ | $8.984 \times 10^0$ |
| 7 | 1993–2023 | 0.03047 | $2.856 \times 10^{-12}$ | $4.154 \times 10^{-6}$ | $9.455 \times 10^2$ |

We then constructed an extended model that simultaneously incorporated all three functional relations:

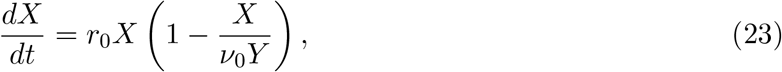

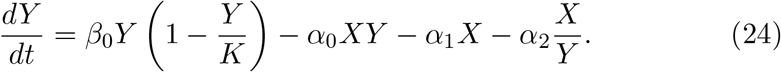

The piecewise-constant framework for this model used six breakpoints and a multi-seed initialization strategy. The extended model produced the best statistical fit among all models. Figure 2 compares observed and model trajectories for *X*(*t*) and *Y* (*t*).

**Figure 2:**
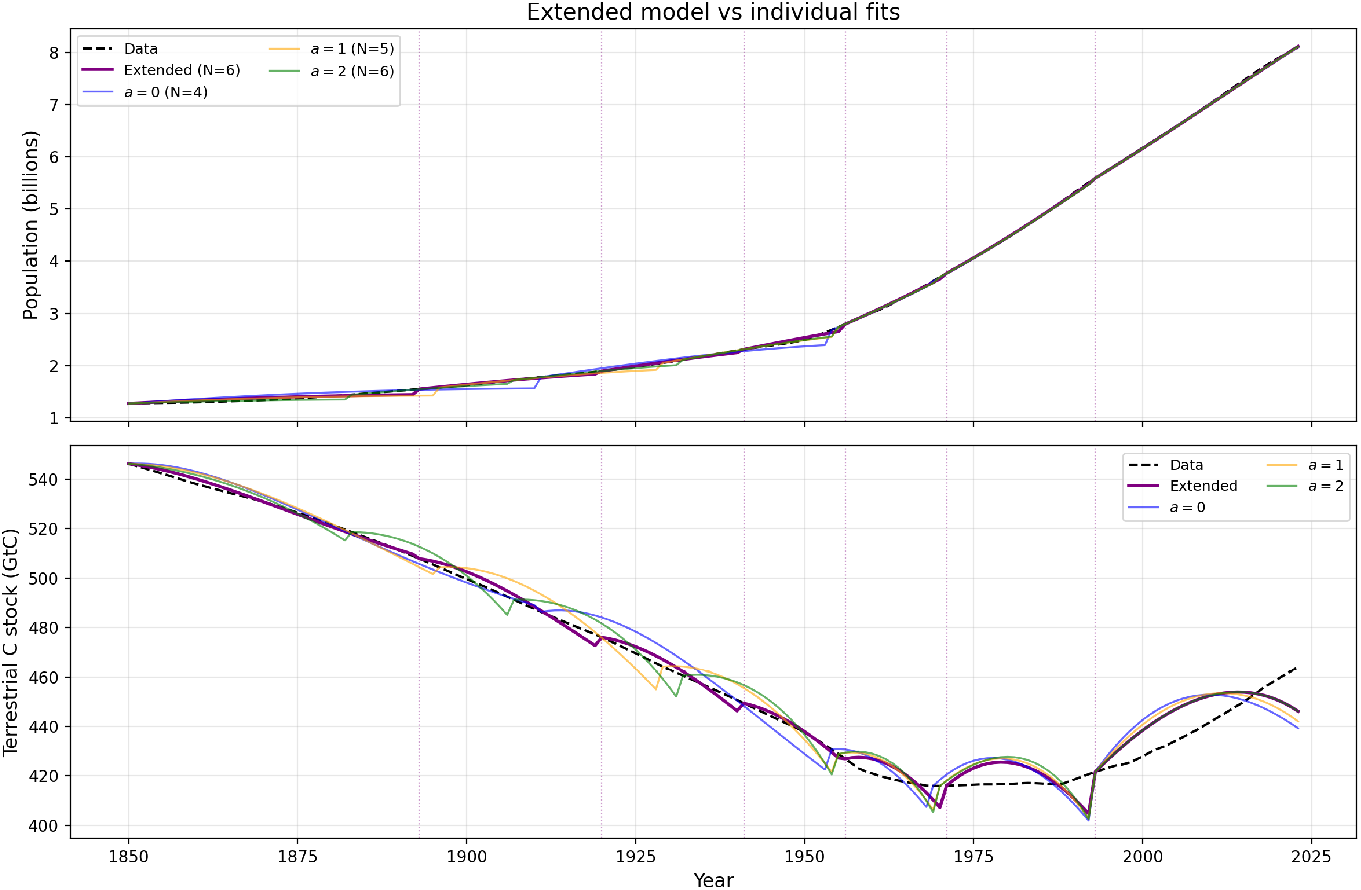
Best fit of the extended model (*N*_bp_ = 6). Observed data (dashed) and model trajectories (solid) for *X*(*t*) and *Y* (*t*).

The phase portraits of the extended model (Figure 3) show the stable existence of a biologically feasible equilibrium embedded in a stable spiral attractor. They also show that the coexistence equilibrium changes through time, increasing in human population size while reducing biosphere stock, until it eventually ceases to exist as a biologically feasible solution. The spiral oscillations also widen, with the current biosphere–human trajectory following the upper-right part of the spiral, associated with strong biosphere depletion together with increasing human population. This trajectory eventually leads to an abrupt population collapse when entering the lower part of the spiral.

**Figure 3:**
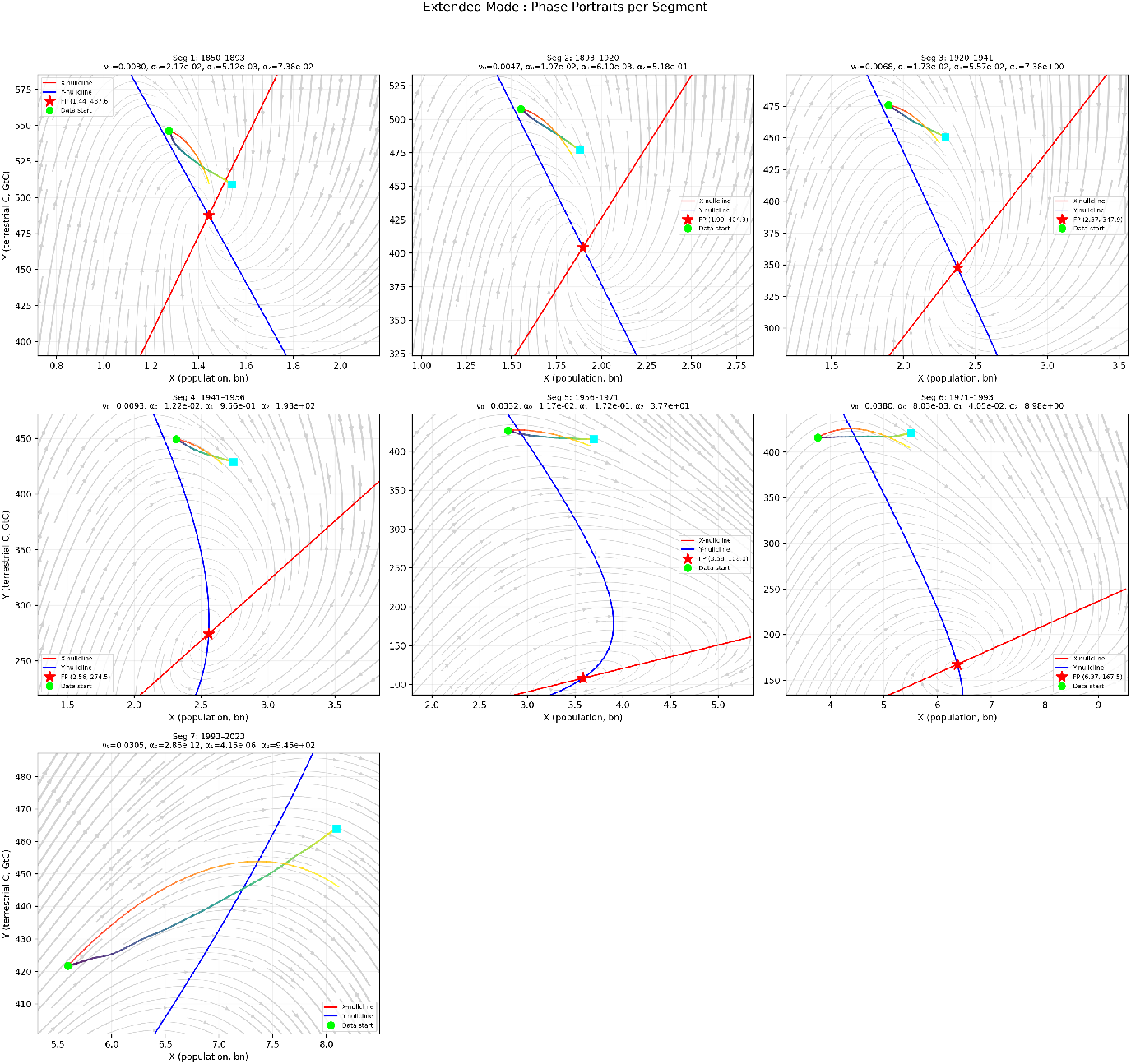
Phase portraits for the extended model across seven segments (*N*_bp_ = 6), showing vector fields, nullclines, coexistence fixed points, observed data, and model trajectories.

We noticed how parameter estimation for *α*’s in the extended model suggests that the predatory-prey functional relation was dominant for most of the time segments, showing greater *consumption shares* in all time segments but the last one (i.e. *α*_0_ weights are greater respect to weights estimated for *α*_1_ and *α*_2_, see supplementary information for share’s calculation), and also, how *ν*_0_ midly increased up to 1950, where it showed a great increase and further stagnation (see Figure 4).

**Figure 4:**
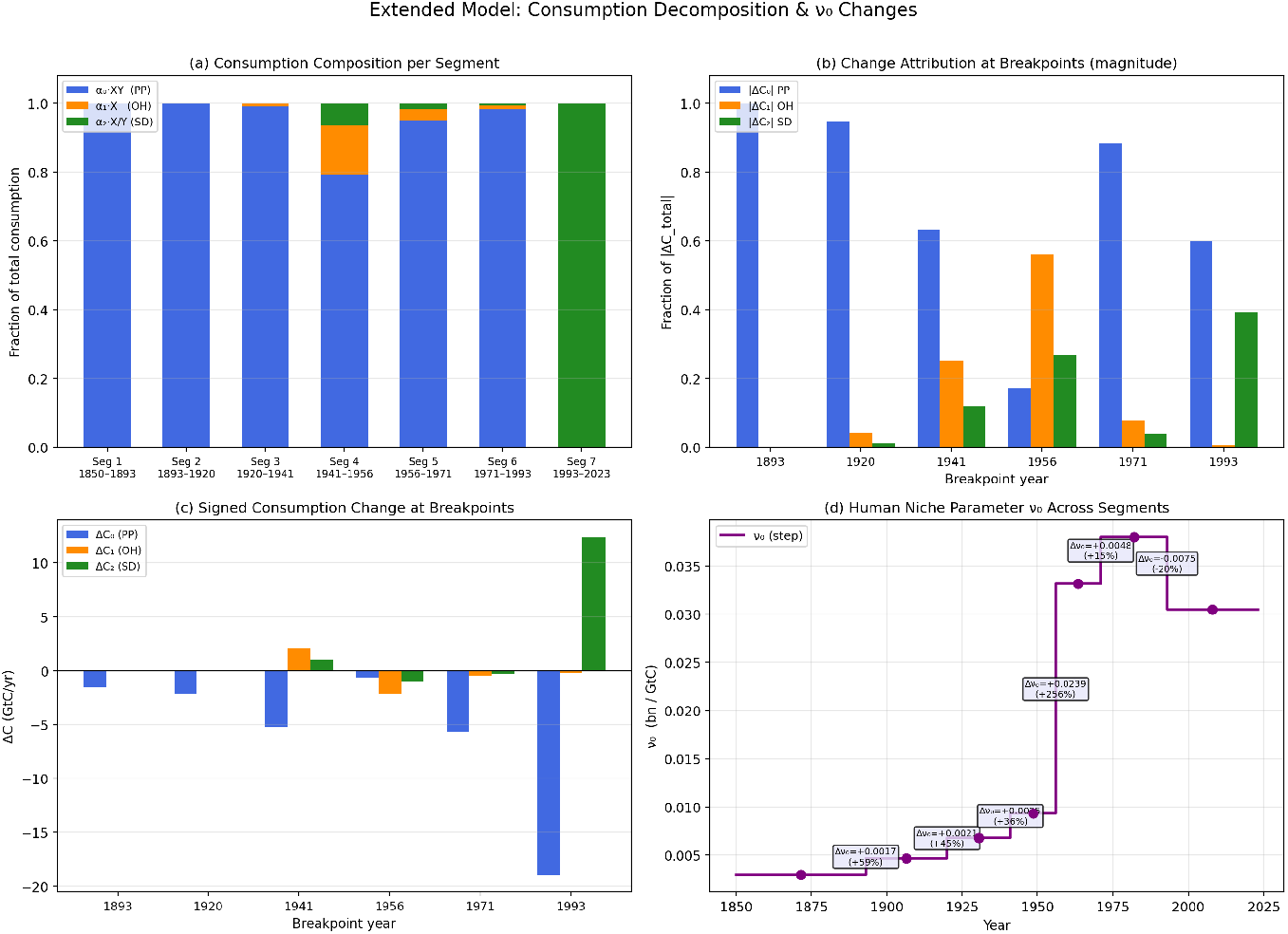
Bootstrap analysis of the extended model (*B* = 300 replicates). **(a)** PP/OH/SD consumption-share distributions by segment (2.5th–97.5th percentile); red stars mark the original-fit shares at segment-mean 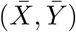. **(b)** *ν*_0_ by segment; stars mark the original fit. **(c)** RSS across replicates; dashed vertical line: original RSS. **(d)** Total effective consumption by segment; stars mark the original fit.

In order to detect the significance of these result, we performed a bootstrap analysis, were we sampled 300 replicates of the model’s residuals and recalculated the relative weights that each *α*’s terms and *ν*_0_ term obtained for all segments. Figure 5 summarizes bootstrap uncertainty for parameters and consumption shares by segment. Here, we confirm how predatory-prey functional relation is dominant for most of the time segments, with more than 50% of replicates selecting greater weights for this relation than for the others. However, in the last innovation period (1993–2023), the dynamics change, with more than 70% of replicates selecting a greater weight on the supply–demand relation, while the predator–prey relation drops to less than 20%. This analysis gives confidence to determine that our predatory relation with the biosphere has changed in the last decades.

**Figure 5:**
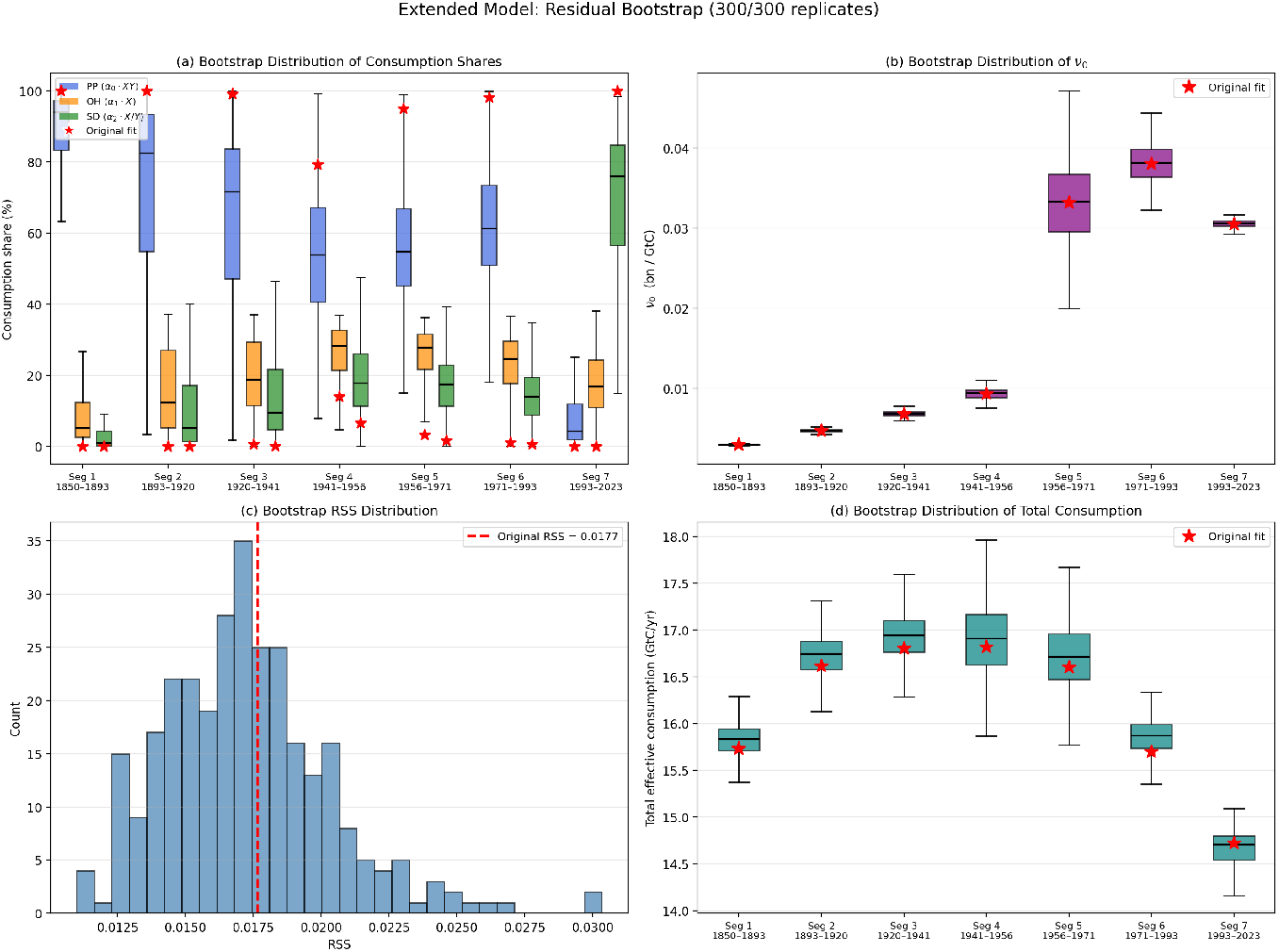
Bootstrap analysis of the extended model (*B* = 300 replicates). **(a)** PP/OH/SD consumption-share distributions by segment (2.5th–97.5th percentile); red stars mark the original-fit shares at segment-mean 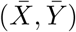. **(b)** *ν*_0_ by segment; stars mark the original fit. **(c)** RSS across replicates; dashed vertical line: original RSS. **(d)** Total effective consumption by segment; stars mark the original fit.

## 4 Discussion

It is interesting that these simple consumer–resource models fit human– terrestrial organic carbon data reasonably well. It is also notable that breakpoints along the historical trajectory correlate with major socioenvironmental changes in human societies and are broadly consistent across models (see Box 1).

**Box 1: Socioenvironmental key changes associated to the 1850-present time-lapse**

**1850–1893**

This time interval relates to the shift from a biomass-to a coal-energy society [Fischer-Kowalski et al., 2014, Smil, 2018]. This transition was characterized by high political instability and armed conflicts [Hobsbawm, 2010, Fischer-Kowalski et al., 2014, Fischer-Kowalski et al., 2019, Fischer-Kowalski et al., 2023]. The energy change from biomass to coal transformed the structure and dynamics of nineteenth-century societies [Fischer-Kowalski, 2011, Fischer-Kowalski et al., 2014]. This shift was gradual, led by the United Kingdom in the first half of the nineteenth century [Smil, 2010]. During the energetic transition from 1860 to 1940, coal emerged as the dominant energy source. The United Kingdom’s share of coal was over 90% from 1850 to 1900, driving substantial growth in energy production [Warde, 2007]. Following this, the United States, France, and Germany also turned to coal as their primary energy source by 1880.

**1893–1920 and 1920–1941**

In 1917, coal accounted for 77% of Japan’s total primary energy; by 1930, it comprised 50% of Russia’s energy consumption [Smil, 2010]. This increase in energy fueled a new era of urbanization and large-scale manufacturing in Great Britain, the United States, Germany, and other coal-producing regions [Mitchell, 2009]. During this period, substantial advances were made in the power and efficiency of steam engines, aligning with the observed increase in energy production rates [Smil, 2018]. After 1900, energy growth rates stagnated, reflecting population and economic trends. The shift from coal to oil and natural gas was accompanied by economic and global financial crises, political instability, and two world wars [Fischer-Kowalski et al., 2019, Fischer-Kowalski et al., 2023]. According to [Mitchell, 2009], the coal era was crucial for understanding the new connections between energy, sociopolitical institutions, mass political movements, and democracy.

**1941–1956, 1956–1971, and 1971–1993**

The second breakpoint occurred around 1940, at the beginning of an era characterized by the rapid expansion of oil and natural gas, known as the “Great Acceleration” (1940–1970; [Steffen et al., 2015]). The shift toward reliance on oil and natural gas [Fischer-Kowalski et al., 2014, Hall and Klitgaard, 2018] caused major upheavals in energy, population, and the economy after 1940–1950. Over the following decades, there was a notable increase in energy, population, and economic growth rates. In the oil era, the energy growth rate tripled, leading to unprecedented levels of material production, population growth, and environmental degradation [Steffen et al., 2015]. The oil and natural gas era transformed social, political, and economic relations inherited from the coal age [Mitchell, 2009]. After 1950, a new energy era emerged, characterized by accelerated population and economic growth and increased human environmental impacts worldwide [Steffen et al., 2015].

**After 1970**

The results showed that from 1970 onward, the dynamics of growth rates in energy, population, and the economy were characterized by diminishing returns [Lima and Berryman, 2011, Hall et al., 2014, Taylor and Tainter, 2016, Bonaiuti, 2018, Lima et al., 2023]. Since 1970, energy, population, and economic growth rates have all decreased; the positive feedback loop among energy, population, and the economy is leading our civilization into a new era of stagnation [Bonaiuti, 2018, Lima et al., 2023]. This trend is closely associated with the observation that the energy return on investment (EROI) of the most important energy sources, such as oil and natural gas, has generally declined over the past fifty years [Hall et al., 2014, Hall and Klitgaard, 2018, Laherrère et al., 2022].

The continuous increase in the technology-modulated parameter *ν*_0_ until around 1950, followed by stagnation, can be interpreted as an initial technological investment in demography-related factors such as sanitation, health, and life expectancy, up to the Great Acceleration, after which the cost of maintaining an individual in modern society increased and fertility rates declined.

Model analyses suggest that we are still on a relatively stable trajectory, in the sense that the system appears to occupy a stable spiral regime. However, the biosphere is being pushed toward lower stocks, and oscillatory dynamics appear to be amplifying. Regular predator–prey dynamics do not show a threat of sudden instability through Hopf bifurcations, but the apparent shift toward a supply–demand interaction is more concerning, because this regime does allow Hopf bifurcations and coexistence instability. New technological relations that preserve both the biosphere and human populations within sustainable trajectories are therefore urgently needed.

## 5 Conclusions

Human predatory consumption of the biosphere can still be sustainable. Because of human agency and sociocultural niche-construction dynamics, it is still possible to innovate in the technology-modulated parameters that regulate sustainable coexistence. Our species has the capacity to occupy a region of parameter space in which both the biosphere and human populations can coexist stably. These models suggest that stable predatory dynamics are still possible, but only if the technological relation between growth and consumption is transformed.

## 6 Open Research

Data and code are freely available in the GitHub repository https://github.com/vanewi/HumansAsPredators. We declare the use of Artifical Intelligence in assisting script coding, through the use of “Cursor” Program. This tool was used for assisting in coding for the model fitting, the bootstrap analysis, and the phase portraits. No IA was used for paper’s conceptualization, data parametrization or results interpretation.

## Acknowledgments

ML was supported by FONDECYT Project #1230075. The authors declare no conflicts of interest.

