## Supplementary Information for "Humans as Predators of the Biosphere: Technological Modulation of Consumer–Resource Dynamics and Its Implications for Sustainability"

#### 1 The General Model: Equilibria and Stability Analyses

The general model follows regular consumer-resource dynamic modeling, with $X$ as the "consumers" (i.e. number of human individuals) and $Y$ the "resource" (i.e. the stock of organic energy or "biosphere's state"). The proposed general human-biosphere model stands as:

$$\frac{dX}{dt} \equiv \dot{X} = r_0 X \left(1 - \frac{X}{\nu_0 Y}\right) \tag{1}$$

$$\frac{dY}{dt} \equiv \dot{Y} = \beta_0 Y \left(1 - \frac{Y}{K}\right) - \alpha_0 X \tag{2}$$

$r_0$ corresponds to the intrinsic human population growth rate; $\beta_0$, the organic energy or biosphere turnover rate; $\nu_0$ is the number of human individuals allocated per unit of organic energy; and $\alpha_0$, the organic energy consumption per human individual, which is modeled through different functional relations, modulated by parameter $a$, as:

$$\alpha_0 X Y^{1-a} = \begin{cases} PREDATOR - PREY: & a = 0 \equiv \alpha_0 Y X \\ ONLY - HUMAN: & a = 1 \equiv \alpha_0 X \\ SUPPLY - DEMAND: & a = 2 \equiv \alpha_0 \frac{X}{Y} \end{cases} \tag{3}$$

We now determine system's equilibria and stability for each particular case.

Table 1 summarizes the model variables and parameters. In particular, the units of $\alpha_0$ depend on the functional relation exponent $a$, since the consumption term $\alpha_0 X Y^{1-a}$ must have units of $[\mathrm{C}]\,[\mathrm{T}]^{-1}$, yielding $[\alpha_0] = [\mathrm{C}]^a\,[\mathrm{ind}]^{-1}\,[\mathrm{T}]^{-1}$.

Table 1: Model variables and parameters. Here "ind" denotes human individuals, "C" denotes mass of terrestrial organic carbon (e.g. GtC), and "T" denotes time (e.g. years).

| Symbol | Description | Units |
|---|---|---|
| *State variables* | | |
| $X$ | Human population | ind |
| $Y$ | Terrestrial organic carbon stock | C |
| $t$ | Time | T |
| *Dimensional parameters* | | |
| $r_0$ | Intrinsic human population growth rate | $T^{-1}$ |
| $\beta_0$ | Biosphere (organic carbon) turnover rate | $T^{-1}$ |
| $K$ | Biosphere carrying capacity | C |
| $\nu_0$ | Human allocation per unit of organic carbon | ind $C^{-1}$ |
| $\alpha_0$ $(a = 0)$ | Per-capita consumption rate (predator-prey) | $\text{ind}^{-1}$ $T^{-1}$ |
| $\alpha_0$ $(a = 1)$ | Per-capita consumption rate (only-human) | C $\text{ind}^{-1}$ $T^{-1}$ |
| $\alpha_0$ $(a = 2)$ | Per-capita consumption rate (supply-demand) | $C^2$ $\text{ind}^{-1}$ $T^{-1}$ |
| *Dimensionless quantities* | | |
| $Y' = Y/K$ | Normalized biosphere state | – |
| $\tau = r_0 t$ | Dimensionless time | – |
| $\beta = \beta_0/r_0$ | Relative biosphere-to-human growth rate | – |
| $\epsilon = \alpha\nu/\beta$ | Compound Human Pressure Index (see §1.1–1.3) | – |

##### 1.1 Case a=0. Predator-Prey Functional Relation

Assuming a mean-field predator-prey functional relation ($a = 0$), the general model stands as:

$$\frac{dX}{dt} \equiv \dot{X} = r_0 X \left(1 - \frac{X}{\nu_0 Y}\right) \tag{4}$$

$$\frac{dY}{dt} \equiv \dot{Y} = \beta_0 Y \left(1 - \frac{Y}{K}\right) - \alpha_0 XY \tag{5}$$

For adimensionalization, let's substitute

$Y'(t) = Y(t)/K$; $\nu = K\nu_0$; $\tau = t * r_0$; $\beta = \beta_0/r_0$; and $\alpha = \alpha_0/r_0$.

The system rewrites to:

$$\frac{dX}{d\tau} \equiv \dot{X} = X \left(1 - \frac{X}{\nu Y'}\right) \tag{6}$$

$$\frac{dY'}{d\tau} \equiv \dot{Y'} = \beta Y' \left(1 - Y'\right) - \alpha XY' \tag{7}$$

System's equilibria are:

$$\begin{cases} trivial, only - biosphere: & X^* = 0,\ Y'^* = 1 \\ trivial, extinction: & X^* = 0,\ Y'^* = 0 \\ non - trivial, coexistence: & X^* = \frac{\nu}{\epsilon+1},\ Y'^* = \frac{1}{\epsilon+1} \end{cases} \tag{8}$$

with $\epsilon = \frac{\alpha\nu}{\beta}$ the *Compound Human Pressure Index*, an adimensional parameter that represents the compound impact of both human population growth capacity and human consumption, normalized by biosphere growth. It is interesting to notice that the value of the non-trivial equilibrium ultimately depends on this parameter.

For analyzing system's stability, we first find the generic Jacobian Matrix:

$$\begin{bmatrix} \frac{\nu Y' - 2X}{\nu Y'} & \frac{X^2}{\nu Y'^2} \\ -\alpha Y' & \beta - 2\beta Y' - \alpha X \end{bmatrix} \tag{9}$$

Now, evaluating the stability for each equilibria:

(i) *Trivial equilibrium for only-biosphere* ($X^* = 0,\ Y'^* = 1$), there are one negative and one positive eigenvalues ($eig = [1,\ -\beta]$), corresponding to an unstable saddle point.

(ii) *Trivial equilibrium for extinction* ($X^* = 0,\ Y'^* = 0$), this equilibrium is of care, because of singularities that arise when $Y'^* = 0$ that have been discussed before (Kot 2011, Safuan et al. 2012).

(iii) *Non-trivial equilibrium* ($X^* = \frac{\nu}{\epsilon+1},\ Y'^* = \frac{1}{\epsilon+1}$). The characteristic polynomial is:

$$\lambda^2 + \frac{\epsilon + \beta + 1}{\epsilon + 1}\,\lambda + \beta = 0 \tag{10}$$

with eigenvalues:

$$\lambda = \frac{-\left(\epsilon + \beta + 1\right) \pm \sqrt{\Delta}}{2\left(\epsilon + 1\right)} \tag{11}$$

where $\Delta = -\left(4\beta - 1\right)\epsilon^2 - 2\epsilon(3\beta - 1) + (\beta - 1)^2$ is the discriminant.

##### 1.1.1 Stability conditions

The **determinant** of the Jacobian at this equilibrium is $\det(J) = \beta > 0$ (since all original rates are positive), so the equilibrium is never a saddle point.

The **trace** is $\mathrm{tr}(J) = -\frac{\epsilon+\beta+1}{\epsilon+1} < 0$ for all $\epsilon > 0,\ \beta > 0$, so the real part of both eigenvalues is always negative. This means:

- The non-trivial equilibrium is **always stable** for any positive parameter values.
- There is **no Hopf bifurcation** (the trace can never change sign), and therefore **no limit cycles** arise from this equilibrium.

##### 1.1.2 Node vs. spiral classification

The transition between stable node (monotonic approach) and stable spiral (damped oscillations) is determined by the sign of the discriminant $\Delta$. Setting $\Delta = 0$ and solving for $\epsilon$ yields the critical curve:

$$\epsilon_c = \frac{\left(\sqrt{\beta}-1\right)^2}{2\sqrt{\beta}-1}, \quad \text{valid for } \beta > \frac{1}{4} \tag{12}$$

The full classification is:

- **For** $0 < \beta \leq \frac{1}{4}$: The discriminant $\Delta > 0$ for all $\epsilon > 0$. The equilibrium is a **stable node** (real negative eigenvalues, monotonic convergence) regardless of the human pressure $\epsilon$.
- **For** $\beta > \frac{1}{4}$:
  - $0 < \epsilon < \epsilon_c$: **Stable node** ($\Delta > 0$, real negative eigenvalues).
  - $\epsilon = \epsilon_c$: **Degenerate stable node** ($\Delta = 0$, repeated negative eigenvalue).
  - $\epsilon > \epsilon_c$: **Stable spiral** ($\Delta < 0$, complex eigenvalues with negative real part; damped oscillations).
- At $\beta = 1$ (i.e. $\beta_0 = r_0$, biosphere and human growth rates are equal), $\epsilon_c = 0$, so the equilibrium is a **stable spiral for all** $\epsilon > 0$.

##### 1.1.3 Dimensional interpretation

The dimensionless parameters in terms of the original (dimensional) quantities are:

$$\beta = \frac{\beta_0}{r_0}, \qquad \epsilon = \frac{\alpha_0\, K\, \nu_0}{\beta_0} \tag{13}$$

where $\beta$ is the ratio of biosphere turnover rate to human growth rate, and $\epsilon$ is the *Compound Human Pressure Index*, measuring the product of per-capita consumption $\alpha_0$, carrying capacity $K$, and human allocation density $\nu_0$, normalized by the biosphere recovery rate $\beta_0$.

#### 1.2 Case a=1. Only-Humans Dynamic

In this case, the functional relation only considers humans ($a = 1$), and the general model stands as:

$$\frac{dX}{dt} \equiv \dot{X} = r_0 X \left(1 - \frac{X}{\nu_0 Y}\right) \tag{14}$$

$$\frac{dY}{dt} \equiv \dot{Y} = \beta_0 Y \left(1 - \frac{Y}{K}\right) - \alpha_0 X \tag{15}$$

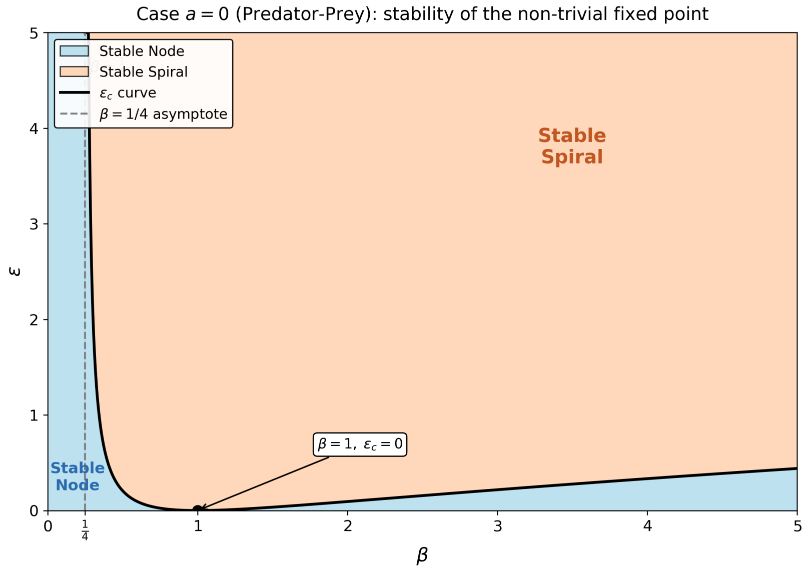


Figure 1: Stability diagram for the non-trivial fixed point in case $a = 0$ (Predator-Prey) in the $(\beta, \epsilon)$ parameter space. The critical curve $\epsilon_c(\beta)$ separates stable nodes from stable spirals. No Hopf bifurcation or instability occurs in this case.

For adimensionalization, let's substitute

$Y'(t) = Y(t)/K$; $\nu = K\nu_0$; $\tau = t * r_0$; $\beta = \beta_0/r_0$; and $\alpha = \frac{\alpha_0}{K\, r_0}$.

The system rewrites to:

$$\frac{dX}{d\tau} \equiv \dot{X} = X\left(1 - \frac{X}{\nu Y'}\right) \tag{16}$$

$$\frac{dY'}{d\tau} \equiv \dot{Y'} = \beta Y'\left(1 - Y'\right) - \alpha X \tag{17}$$

System's equilibria are:

$$\begin{cases} trivial, only-biosphere: & X^* = 0,\ Y'^* = 1 \\ trivial, extinction: & X^* = 0,\ Y'^* = 0 \\ non-trivial, coexistence: & X^* = \nu\left(1-\epsilon\right),\ Y'^* = 1-\epsilon \end{cases} \tag{18}$$

It is required that $\epsilon < 1$ for the equilibria to be biologically relevant (not negative).

For analyzing system's stability, we first find the generic Jacobian Matrix:

$$\begin{bmatrix} \frac{\nu Y' - 2X}{\nu Y'} & \frac{X^2}{\nu Y'^2} \\ -\alpha & \beta - 2\beta Y' \end{bmatrix} \tag{19}$$

Now, evaluating the stability for each equilibria:

(i) *Trivial equilibrium for only-biosphere* ($X^* = 0,\ Y'^* = 1$), stands the same as before, with one negative and one positive eigenvalues ($eig = [1,\ -\beta]$), corresponding to an unstable saddle point.

(ii) *Trivial equilibrium for extinction* ($X^* = 0,\ Y'^* = 0$), stands the same as before, with singularities arising when $Y'^* = 0$ (Kot 2011, Safuan et al. 2012).

(iii) *Non-trivial equilibrium* ($X^* = \nu\left(1-\epsilon\right),\ Y'^* = 1-\epsilon$, valid for $0 < \epsilon < 1$). The characteristic polynomial is:

$$\lambda^2 - (2\beta\epsilon - \beta - 1)\,\lambda + \beta(1-\epsilon) = 0 \tag{20}$$

with eigenvalues:

$$\lambda = \frac{(2\beta\epsilon - \beta - 1) \pm \sqrt{\Delta}}{2} \tag{21}$$

where $\Delta = 4\beta^2\epsilon^2 - 4\beta^2\epsilon + (\beta-1)^2$ is the discriminant.

##### 1.2.1 Stability conditions

The **determinant** is $\det(J) = \beta(1-\epsilon) > 0$ for $\epsilon \in (0,1)$, so the equilibrium is never a saddle point within its existence range. At $\epsilon = 1$, $\det(J) = 0$ and the non-trivial equilibrium merges with the extinction equilibrium (transcritical bifurcation).

The **trace** is $\mathrm{tr}(J) = 2\beta\epsilon - \beta - 1$, which — unlike case $a = 0$ — can change sign. Setting $\mathrm{tr}(J) = 0$ yields a **Hopf bifurcation** threshold at:

$$\epsilon_H = \frac{\beta+1}{2\beta} \tag{22}$$

This threshold lies in the valid range $(0,1)$ only when $\beta > 1$. The transversality condition is satisfied since $\frac{d}{d\epsilon}\mathrm{tr}(J) = 2\beta > 0$.

- **For** $0 < \beta \leq 1$: $\epsilon_H \geq 1$, so $\mathrm{tr}(J) < 0$ for all $\epsilon \in (0,1)$. The equilibrium is **always stable**; no Hopf bifurcation occurs.
- **For** $\beta > 1$: $\epsilon_H \in \left(\frac{1}{2}, 1\right)$, and:
  - $0 < \epsilon < \epsilon_H$: $\mathrm{tr}(J) < 0 \Rightarrow$ **stable**.
  - $\epsilon = \epsilon_H$: $\mathrm{tr}(J) = 0 \Rightarrow$ **Hopf bifurcation** (purely imaginary eigenvalues $\lambda = \pm i\sqrt{\frac{\beta-1}{2}}$).
  - $\epsilon_H < \epsilon < 1$: $\mathrm{tr}(J) > 0 \Rightarrow$ **unstable**.

##### 1.2.2 Node vs. spiral classification

Setting $\Delta = 0$ and solving for $\epsilon$ yields:

$$\epsilon_{c\pm} = \frac{\beta \pm \sqrt{2\beta - 1}}{2\beta}, \quad \text{valid for } \beta \geq \frac{1}{2} \tag{23}$$

Both roots satisfy $0 < \epsilon_{c-} < \epsilon_{c+} < 1$ for $\beta > 1/2$ and $\beta \neq 1$. Since the leading coefficient of $\Delta$ (as a quadratic in $\epsilon$) is $4\beta^2 > 0$, we have $\Delta < 0$

(spiral) for $\epsilon_{c-} < \epsilon < \epsilon_{c+}$ and $\Delta > 0$ (node) otherwise. When $\beta > 1$, one can verify that $\epsilon_{c-} < \epsilon_H < \epsilon_{c+}$, confirming that the Hopf bifurcation occurs within the spiral region.

The full classification is:

- **For** $0 < \beta < \frac{1}{2}$: **Stable node** for all $\epsilon \in (0, 1)$.
- **For** $\frac{1}{2} \leq \beta \leq 1$ (always stable, no Hopf):
  - $0 < \epsilon < \epsilon_{c-}$: **Stable node**.
  - $\epsilon_{c-} < \epsilon < \epsilon_{c+}$: **Stable spiral** (damped oscillations).
  - $\epsilon_{c+} < \epsilon < 1$: **Stable node**.
- **For** $\beta > 1$ (Hopf bifurcation at $\epsilon_H$):
  - $0 < \epsilon < \epsilon_{c-}$: **Stable node**.
  - $\epsilon_{c-} < \epsilon < \epsilon_H$: **Stable spiral**.
  - $\epsilon = \epsilon_H$: **Center** (Hopf bifurcation; possible limit cycle birth).
  - $\epsilon_H < \epsilon < \epsilon_{c+}$: **Unstable spiral**.
  - $\epsilon_{c+} < \epsilon < 1$: **Unstable node**.
- At $\beta = 1$: $\epsilon_{c-} = 0$ and $\epsilon_{c+} = 1$, so the equilibrium is a **stable spiral for all** $\epsilon \in (0, 1)$. The Hopf threshold $\epsilon_H = 1$ coincides with the transcritical bifurcation, so no instability arises.

##### 1.2.3 Dimensional interpretation

The dimensionless parameters in terms of the original (dimensional) quantities are:

$$\beta = \frac{\beta_0}{r_0}, \qquad \epsilon = \frac{\alpha_0 \, \nu_0}{\beta_0} \tag{24}$$

Compared to case $a = 0$, the carrying capacity $K$ no longer appears in $\epsilon$. This reflects the fact that in the $a = 1$ interaction the consumption term $\alpha_0 X$ does not depend on $Y$, so the biosphere's size only enters through the nondimensionalization of $Y$ itself, not through the effective pressure index.

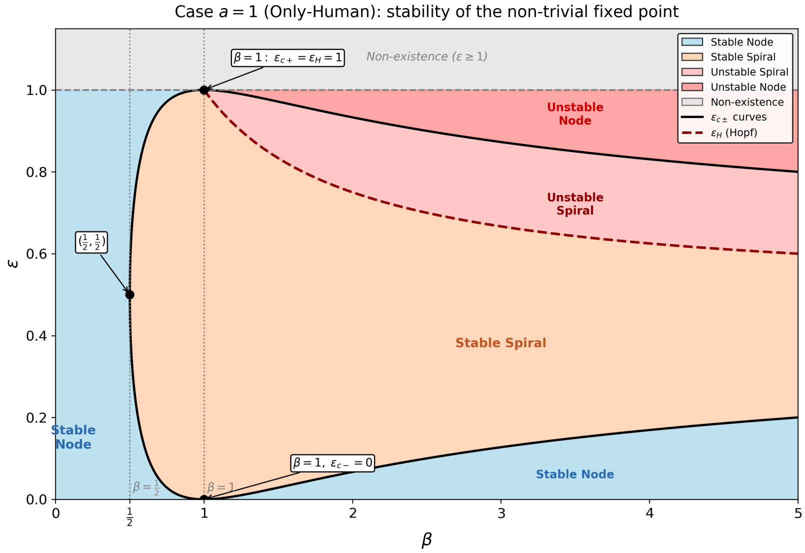


Figure 2: Stability diagram for the non-trivial fixed point in case $a = 1$ (Only-Human) in the $(\beta, \epsilon)$ parameter space. The discriminant curves $\epsilon_{c\pm}(\beta)$ separate nodes from spirals, while the Hopf line $\epsilon_H(\beta)$ separates stable from unstable dynamics. The non-trivial equilibrium exists only for $\epsilon < 1$ (transcritical bifurcation).

##### 1.3 Case a=2. Supply-Demand Dynamic

In this case ($a = 2$), the functional relation depends on the ratio of humans $X$ vs. biospheric state $Y$, and the general model stands as:

$$\frac{dX}{dt} \equiv \dot{X} = r_0 X \left(1 - \frac{X}{\nu_0 Y}\right) \tag{25}$$

$$\frac{dY}{dt} \equiv \dot{Y} = \beta_0 Y \left(1 - \frac{Y}{K}\right) - \alpha_0 \frac{X}{Y} \tag{26}$$

For adimensionalization, let's substitute

$Y'(t) = Y(t)/K$; $\nu = K\nu_0$; $\tau = t * r_0$; $\beta = \beta_0/r_0$; and $\alpha = \frac{\alpha_0}{K^2 r_0}$.

The system rewrites to:

$$\frac{dX}{d\tau} \equiv \dot{X} = X \left(1 - \frac{X}{\nu Y'}\right) \tag{27}$$

$$\frac{dY'}{d\tau} \equiv \dot{Y'} = \beta Y' \left(1 - Y'\right) - \alpha \frac{X}{Y'} \tag{28}$$

System's equilibria are:

$$\begin{cases} trivial, only - biosphere: & X^* = 0,\ Y'^* = 1 \\ trivial, extinction: & X^* = 0,\ Y'^* = 0 \\ non - trivial, coexistence: & X^* = \frac{\nu\left(1 \pm \sqrt{1-4\epsilon}\right)}{2},\ Y'^* = \frac{1 \pm \sqrt{1-4\epsilon}}{2} \end{cases} \tag{29}$$

Notice that different non-trivial equilibria can exist depending on $\epsilon$: for $\epsilon < \frac{1}{4}$, two distinct coexistence equilibria exist; for $\epsilon = \frac{1}{4}$, a single equilibrium at $Y'^* = \frac{1}{2}$; and for $\epsilon > \frac{1}{4}$, no real coexistence equilibrium exists.

For analyzing system's stability, we first find the generic Jacobian Matrix:

$$\begin{bmatrix} \frac{\nu Y' - 2X}{\nu Y'} & \frac{X^2}{\nu Y'^2} \\ -\frac{\alpha}{Y'} & -\frac{2\beta Y'^3 - \beta Y'^2 - \alpha X}{Y'^2} \end{bmatrix} \tag{30}$$

Now, evaluating the stability for each equilibria:

(i) *Trivial equilibrium for only-biosphere* ($X^* = 0,\ Y'^* = 1$), stands the same as before, with one negative and one positive eigenvalues ($eig = [1,\ -\beta]$), corresponding to an unstable saddle point.

(ii) *Trivial equilibrium for extinction* ($X^* = 0,\ Y'^* = 0$), stands the same as before, with singularities arising when $Y'^* = 0$ (Kot 2011, Safuan et al. 2012).

(iii) *Non-trivial equilibria* ($X^* = \nu Y'^*$, with $Y'^*_{\pm} = \frac{1 \pm \sqrt{1-4\epsilon}}{2}$, existing for $0 < \epsilon < \frac{1}{4}$). At $\epsilon = \frac{1}{4}$ the two equilibria merge at $Y'^* = \frac{1}{2}$ and annihilate (saddle-node bifurcation).

Evaluating the Jacobian at $X^* = \nu Y'^*$, and using the equilibrium condition $\alpha\nu = \beta Y'^*(1 - Y'^*)$, the trace and determinant simplify to:

$$\mathrm{tr}(J) = 2\beta - 3\beta Y'^* - 1 \tag{31}$$

$$\det(J) = \beta\left(2Y'^* - 1\right) \tag{32}$$

yielding the characteristic polynomial:

$$\lambda^2 - (2\beta - 3\beta Y'^* - 1)\,\lambda + \beta(2Y'^* - 1) = 0 \tag{33}$$

with eigenvalues:

$$\lambda = \frac{(2\beta - 3\beta Y'^* - 1) \pm \sqrt{\Delta}}{2} \tag{34}$$

where $\Delta = 9\beta^2 Y'^{*2} - 2\beta(6\beta + 1)Y'^* + 4\beta^2 + 1$ is the discriminant.

##### 1.3.1 The lower equilibrium $Y'^*_-$ is always a saddle

For $Y'^*_- = \frac{1-\sqrt{1-4\epsilon}}{2} < \frac{1}{2}$, we have $2Y'^*_- - 1 = -\sqrt{1-4\epsilon} < 0$, so $\det(J) = -\beta\sqrt{1-4\epsilon} < 0$. This means the eigenvalues are real with opposite signs: the lower equilibrium is **always a saddle point**.

##### 1.3.2 Stability of the upper equilibrium $Y'^*_+$

For $Y'^*_+ = \frac{1+\sqrt{1-4\epsilon}}{2} > \frac{1}{2}$, we have $\det(J) = \beta\sqrt{1-4\epsilon} > 0$, so the upper equilibrium is never a saddle.

The trace is $\mathrm{tr}(J) = 2\beta - 3\beta Y'^*_+ - 1$, which is a monotonically increasing function of $\epsilon$ (since $Y'^*_+$ decreases with $\epsilon$). At the endpoints:

- $\epsilon \to 0$ ($Y'^*_+ \to 1$): $\mathrm{tr}(J) \to -\beta - 1 < 0$ (stable).
- $\epsilon \to \frac{1}{4}$ ($Y'^*_+ \to \frac{1}{2}$): $\mathrm{tr}(J) \to \frac{\beta}{2} - 1$, which is negative for $\beta < 2$, zero at $\beta = 2$, and positive for $\beta > 2$.

Setting $\mathrm{tr}(J) = 0$ yields a **Hopf bifurcation** at:

$$\epsilon_H = \frac{(2\beta - 1)(\beta + 1)}{9\beta^2}, \quad \text{valid for } \beta > 2 \tag{35}$$

corresponding to $Y'^*_+ = \frac{2\beta-1}{3\beta}$ and purely imaginary eigenvalues $\lambda = \pm i\sqrt{\frac{\beta-2}{3}}$. The transversality condition is satisfied since $\frac{d}{d\epsilon}\mathrm{tr}(J) = \frac{3\beta}{\sqrt{1-4\epsilon}} > 0$.

For $\beta \leq 2$, the trace remains non-positive throughout the existence range, and the upper equilibrium is **always stable**. At $\beta = 2$, the Hopf threshold $\epsilon_H = \frac{1}{4}$ coincides with the saddle-node bifurcation (a Bogdanov–Takens codimension-2 point).

###### 1.3.3 Node vs. spiral classification

The discriminant evaluated at the equilibrium endpoints gives $\Delta|_{Y'^*=1} = (\beta - 1)^2$ and $\Delta|_{Y'^*=1/2} = \frac{(\beta-2)^2}{4}$, both non-negative. The minimum of $\Delta$ (over $Y'^*$) is $\Delta_{\min} = -\frac{4(3\beta-2)}{9}$, so a spiral region (where $\Delta < 0$) exists only for $\beta > \frac{2}{3}$.

For $\beta > \frac{2}{3}$, the discriminant vanishes at:

$$Y'^*_{d\pm} = \frac{(6\beta + 1) \pm 2\sqrt{3\beta - 2}}{9\beta} \tag{36}$$

with $\frac{1}{2} \leq Y'^*_{d-} \leq Y'^*_{d+} \leq 1$ (equalities at $\beta = 2$ and $\beta = 1$ respectively). The spiral region is $Y'^*_{d-} < Y'^* < Y'^*_{d+}$, and one can verify that for $\beta > 2$, the Hopf point $Y'^*_H = \frac{2\beta-1}{3\beta}$ lies within this spiral region.

###### 1.3.4 Full classification of the upper equilibrium

- **For** $0 < \beta \leq \frac{2}{3}$: $\Delta \geq 0$ and $\mathrm{tr}(J) < 0$ for all $\epsilon \in (0, \frac{1}{4})$. The equilibrium is a **stable node**.
- **For** $\frac{2}{3} < \beta < 2$ (always stable, no Hopf): As $\epsilon$ increases from 0 to $\frac{1}{4}$:
  - **Stable node** $\to$ **Stable spiral** $\to$ **Stable node**.

- **For** $\beta = 2$ (Bogdanov–Takens point): The Hopf threshold $\epsilon_H = \frac{1}{4}$ coincides with the saddle-node bifurcation, so the equilibrium remains stable throughout its existence. As $\epsilon$ increases:
  - **Stable node** $\to$ **Stable spiral** $\to$ degenerate at $\epsilon = \frac{1}{4}$.
- **For** $\beta > 2$ (Hopf bifurcation at $\epsilon_H < \frac{1}{4}$): As $\epsilon$ increases from 0 to $\frac{1}{4}$:
  - **Stable node** $\to$ **Stable spiral** $\to$ **Hopf bifurcation** (possible limit cycle birth) $\to$ **Unstable spiral** $\to$ **Unstable node** $\to$ saddle-node annihilation.

As $\beta \to \infty$, $\epsilon_H \to \frac{2}{9}$ and the unstable window $(\epsilon_H, \frac{1}{4})$ approaches a fixed width.

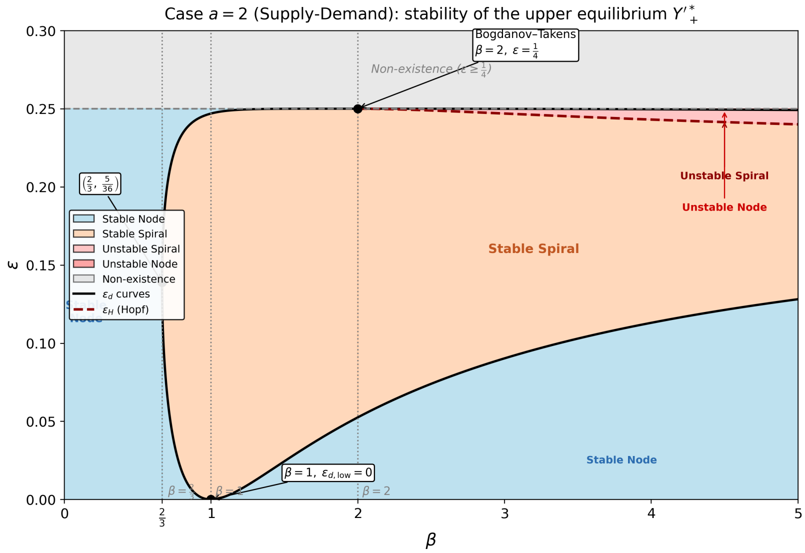


Figure 3: Stability diagram for the upper equilibrium $Y'^*_+$ in case $a = 2$ (Supply-Demand) in the $(\beta, \epsilon)$ parameter space. Both non-trivial equilibria exist only for $\epsilon < \frac{1}{4}$ (saddle-node boundary); the lower equilibrium $Y'^*_-$ is always a saddle (not shown). The Bogdanov–Takens codimension-2 point at $(\beta, \epsilon) = (2, \frac{1}{4})$ organizes the transition between stable and unstable dynamics.

##### 1.3.5 Dimensional interpretation

The dimensionless parameters in terms of the original (dimensional) quantities are:

$$\beta = \frac{\beta_0}{r_0}, \qquad \epsilon = \frac{\alpha_0\, \nu_0}{K\, \beta_0} \tag{37}$$

Compared to the previous cases, $\epsilon$ now includes an additional factor of $K^{-1}$. This reflects the supply-demand nature of the $a = 2$ interaction: because the consumption rate $\alpha_0 X/Y$ is inversely proportional to the biosphere state $Y$, a larger carrying capacity $K$ dilutes the effective human pressure, entering $\epsilon$ in the denominator. Consequently, a richer biosphere (larger $K$) reduces $\epsilon$ and promotes stability, whereas in case $a = 0$ a larger $K$ *increases* $\epsilon$ and pushes the system toward oscillatory dynamics.

### 2 Parameter Estimation from Empirical Data

The dimensionless parameters $\beta = \beta_0/r_0$ and $\epsilon$ (whose dimensional form depends on $a$; see §1.1–1.3) govern the stability of the model's equilibria. Here we estimate the underlying dimensional parameters $K$, $\beta_0$, and $r_0$ from current empirical data, and derive the implied range of $\beta$.

First, several studies on hunter-gatherer populations report basic demographic attributes such as reproductive span, spacing period, and population growth rate (Hassan 1973, Pennington 1996, Gurven et al. 2007). As $\nu_0$ refers to the maximum per-capita growth rate of the human population when resources are abundant and density-dependent limitation is negligible, this parameter could go as high as 0.04yr $^{-1}$. However, we decide to use a conservative baseline of 0.025yr $^{-1}$.

Then, recalling that this study addresses the biosphere as the total organic carbon stock fixed by terrestrial primary producers, we first determined that current census of Earth biomass calculated that plants account for 450 GtC (Bar-On et al. 2018). On another hand, Erb et al. (2018) determined the potential vegetation that would exist under current climate but without any human land use, estimating a $K$ of 916 GtC. Organic carbon turnover rate $\beta_0$ is estimated as the ratio of net primary productivity (NPP) to the standing biomass stock, or the inverse of the turnover time in a system at equilibrium. Estimating NPP has been an active research area for decades, and a conservative value is established at 56 PgCyr $^{-1}$ (Erb et al. 2018, Hoehler et al. 2023). This gives a $\beta_0 = 0.067$ yr$^{-1}$.

The current NPP of $\sim 58\,\mathrm{PgC\,yr^{-1}}$ corresponds to the present-day biosphere at $\sim 450\,\mathrm{GtC}$, already degraded to roughly half of $K$. The estimated $\beta_0$ needs the turnover time of the *natural* (undisturbed) biosphere.

Erb et al. (2016) showed that human land use has accelerated vegetation biomass turnover by a factor of $\sim 1.9$ globally (halving the residence time), with 59% due to land conversion, 26% to forestry, and 15% to grassland use. The current vegetation turnover time is $\tau_{\text{current}} = 450/58 \approx 8\,\text{yr}$, implying a natural turnover time of

$$\tau_{\text{natural}} \approx 1.9 \times 8 \approx 15\ \text{yr}.$$

This is consistent with the whole-ecosystem carbon turnover time of $\sim 23\,\text{yr}$ estimated by Carvalhais et al. (2014) from a global observation-based assessment (the vegetation-only component is shorter than the total, which includes slower soil pools). Therefore:

$$\boxed{\beta_0 \approx \frac{1}{15} \approx 0.067\ \text{yr}^{-1}} \tag{38}$$

As a cross-check, the natural-equilibrium NPP implied by this rate is $\beta_0 K \approx 0.067 \times 916 \approx 61\,\mathrm{PgC\,yr^{-1}}$, consistent with the indirect GPP $\times$ CUE estimate.

Table 2: Key parameter values adopted for the fitted model, together with their empirical basis. The carrying-capacity value keeps the explicit potential-vegetation estimate of 916 GtC from Erb et al. (2018), while $r_0$ uses the central value selected from the empirically motivated range in Cohen (1995).

| **Parameter** | **Value used** | **Reference / basis** |
|---|---|---|
| $K$ | 916 GtC | Erb et al. (2018); Spawn et al. (2020) |
| $\beta_0$ | $0.067\,\text{yr}^{-1}$ | Erb et al. (2016); Endsley et al. (2023) |
| $r_0$ | $0.025\,\text{yr}^{-1}$ | Cohen (1995), central value from 0.02–0.03 $\text{yr}^{-1}$ |
| $\beta = \beta_0/r_0$ | 2.68 | derived from adopted $K$, $\beta_0$, and $r_0$ |

The range $\beta \approx 2$–3 places the system squarely within the dynamically rich regime of the stability diagrams (Figures 1–3):

- In case $a = 0$: always stable, in the spiral region for most $\epsilon$ values.
- In case $a = 1$: $\beta > 1$, so the Hopf bifurcation threshold $\epsilon_H$ is active; the system can become unstable if $\epsilon > \epsilon_H$.

- In case $a = 2$: $\beta > 2$, so the Hopf bifurcation at $\epsilon_H < \frac{1}{4}$ is active and a window of instability exists before the saddle-node annihilation.

### 3 Empirical Data

Two primary data sources underpin the empirical analysis:

- **Human population** $X(t)$: Global population from 1850 to 2023 (in billions of individuals), obtained from Our World in Data (OWID), which compiles historical estimates from the United Nations and the History Database of the Global Environment (HYDE).

- **Land-use-change $CO_2$ emissions**: Annual land-use-change (LUC) $CO_2$ emissions from the Global Carbon Budget (GCB; Friedlingstein et al., 2025), converted from $MtCO_2$ to GtC via the factor 12/44,000.

#### 3.1 Reconstruction of terrestrial carbon stock $Y(t)$

The terrestrial organic carbon stock $Y(t)$ is not directly observed at annual resolution over the full period. We reconstruct it from the net carbon flux into the terrestrial biosphere as follows. Define the net flux as

$$F_{\text{net}}(t) \;=\; S_{\text{land}}(t) \;-\; E_{\text{LUC}}(t), \tag{39}$$

where $S_{\text{land}}(t)$ is the decade-averaged natural land carbon sink (from GCB and TRENDY ensemble estimates in the literature) and $E_{\text{LUC}}(t)$ is the annual LUC emission. The stock is then computed as

$$Y(t) \;=\; Y_{1850} \;+\; \sum_{s=1851}^{t} F_{\text{net}}(s), \tag{40}$$

with the integration constant $Y_{1850}$ chosen so that $Y(2015) \approx 450\,\text{GtC}$, consistent with the biomass census of Bar-On et al. (2018).

Both $X(t)$ and $Y(t)$ are interpolated to a common annual grid of $N_{\text{data}} = 174$ points (1850–2023).

### 4 Piecewise-Constant Parameter Estimation

#### 4.1 Fitting framework

Because the interaction parameters $\nu_0$ and $\alpha_0$ are expected to change over the 174-year record due to technological and socioeconomic shifts, we model

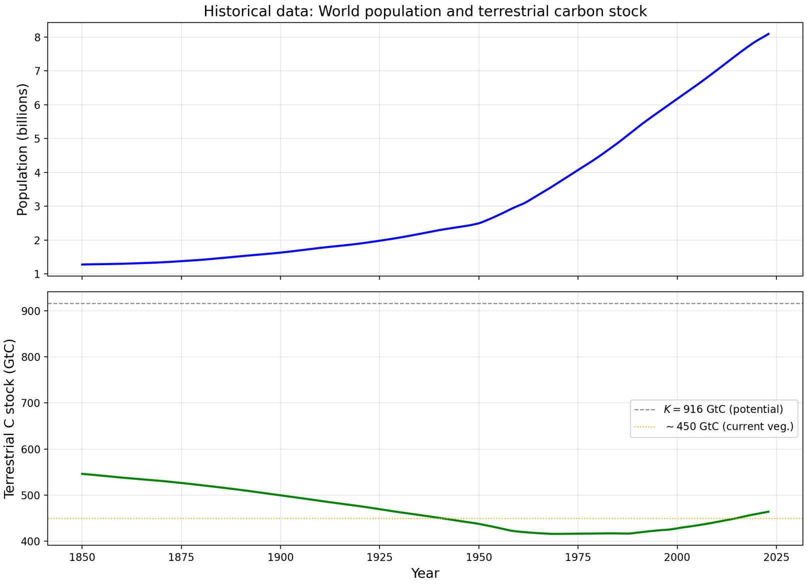


Figure 4: Reconstructed time series of global human population $X(t)$ (left axis, billions) and terrestrial organic carbon stock $Y(t)$ (right axis, GtC) over 1850–2023.

them as piecewise-constant functions over $N_{\rm bp}+1$ segments separated by $N_{\rm bp}$ breakpoints $\{t_{b_1},\ldots,t_{b_{N_{\rm bp}}}\}$. For each individual model ($a=0,1,2$), the number of free parameters is

$$n_{\rm params} = 3\,N_{\rm bp} + 2, \tag{41}$$

corresponding to one value of $\nu_0$ and one of $\alpha_0$ per segment (i.e. $2(N_{\rm bp}+1)$ interaction parameters), plus $N_{\rm bp}$ breakpoint times.

###### 4.1.1 Multiple-shooting formulation

The ODE system is integrated independently on each segment via a multiple-shooting approach: at each breakpoint $t_{b_j}$, the initial conditions for the next segment are re-anchored from the observed data $(X(t_{b_j}),\, Y(t_{b_j}))$. This avoids error accumulation across segments and improves numerical stability.

###### 4.1.2 Model selection via Kneedle/elbow detection

The optimal number of breakpoints $N_{\rm bp}$ is selected using the Kneedle (elbow) algorithm applied to the log(RSS) vs. $N_{\rm bp}$ curve. This model-free geometric approach identifies the point of maximum curvature—i.e., the "knee" where adding further breakpoints yields diminishing returns in fit quality.

Standard information criteria (BIC, AICc) were explored but found unreliable for this problem: the autocorrelation inherent in ODE-generated time series inflates the effective sample size, causing BIC to consistently favor high $N_{\rm bp}$ values ($\geq$16) without converging. The Kneedle approach avoids parametric assumptions about the penalty structure and provides robust, case-specific breakpoint counts.

###### 4.1.3 Optimization procedure

Fitting is implemented in PyTorch with `torchdiffeq` for differentiable ODE integration. Initialization is obtained via gradient matching: Savitzky–Golay smoothing of the observed trajectories followed by dynamic programming to determine optimal breakpoint placement. Parameters are stored in log-space to guarantee positivity. The initial estimates are then refined in two phases: Adam optimizer (learning rate $10^{-2}$, 300 iterations with gradient clipping) followed by L-BFGS quasi-Newton refinement (100 iterations, strong Wolfe line search).

#### 4.2 Individual model results

Table 3 summarizes the best-fit results for each of the three individual models.

Table 3: Best-fit results for each individual model ($a = 0, 1, 2$), with $N_{\text{bp}}$ selected independently per case via Kneedle/elbow detection.

| **Model** | $a$ | $N_{\text{bp}}$ | $n_{\text{params}}$ | RSS |
|---|---|---|---|---|
| Predator–Prey | 0 | 4 | 14 | 0.0414 |
| Only-Human | 1 | 5 | 17 | 0.0313 |
| Supply–Demand | 2 | 6 | 20 | 0.0252 |

The Kneedle algorithm selects different $N_{\text{bp}}$ for each case: 4 for Predator–Prey, 5 for Only-Human, and 6 for Supply–Demand. Despite these differences, the breakpoints cluster near common historical transition dates: the early 1900s industrialization wave, the interwar period, the post-WWII economic expansion, the 1970 energy crisis, and the onset of globalization and digital technology (∼1993).

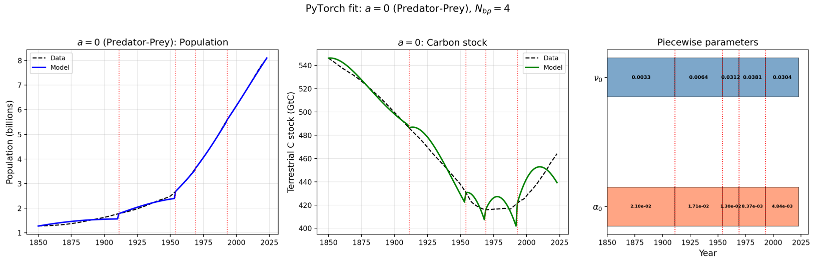


Figure 5: Best fit for the Predator–Prey model ($a = 0$, $N_{\text{bp}} = 4$). Observed data (dashed) and model trajectories (solid) for $X(t)$ and $Y(t)$, with piecewise parameter timeline.

#### 4.3 Epsilon trajectories on stability diagrams

The estimated piecewise-constant parameters allow us to compute the dimensionless Compound Human Pressure Index $\epsilon$ for each segment and overlay these values on the stability diagrams derived in Section 1. Figure 8 shows the resulting trajectories in the $(\beta, \epsilon)$ plane, indicating the dynamical regime occupied by the system during each historical period.

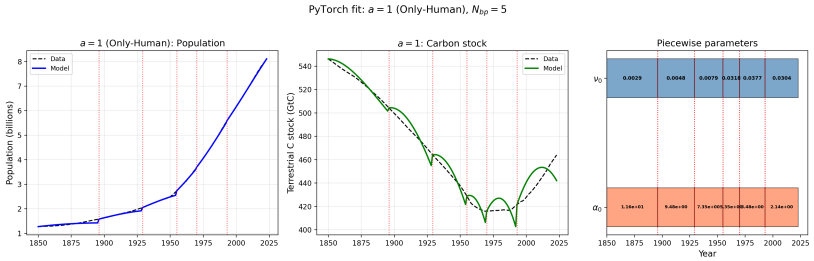


Figure 6: Best fit for the Only-Human model ($a = 1$, $N_{\text{bp}} = 5$). Observed data (dashed) and model trajectories (solid) for $X(t)$ and $Y(t)$, with piecewise parameter timeline.

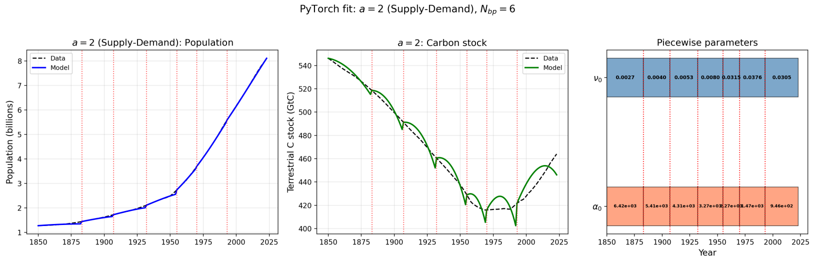


Figure 7: Best fit for the Supply–Demand model ($a = 2$, $N_{\text{bp}} = 6$). Observed data (dashed) and model trajectories (solid) for $X(t)$ and $Y(t)$, with piecewise parameter timeline.

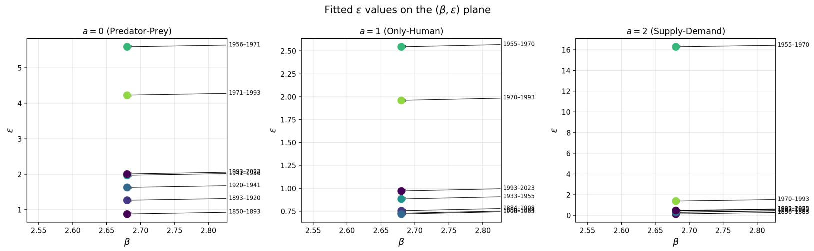


Figure 8: Estimated $\epsilon$ values per segment overlaid on the stability diagrams for each case ($a = 0, 1, 2$). Each point represents one piecewise-constant segment; arrows indicate the chronological progression.

##### 4.4 Phase portraits

Figures 9–11 display phase portraits for each segment of each individual model. Each panel shows the vector field, nullclines, non-trivial fixed points, and both the observed data trajectory and the fitted model trajectory.

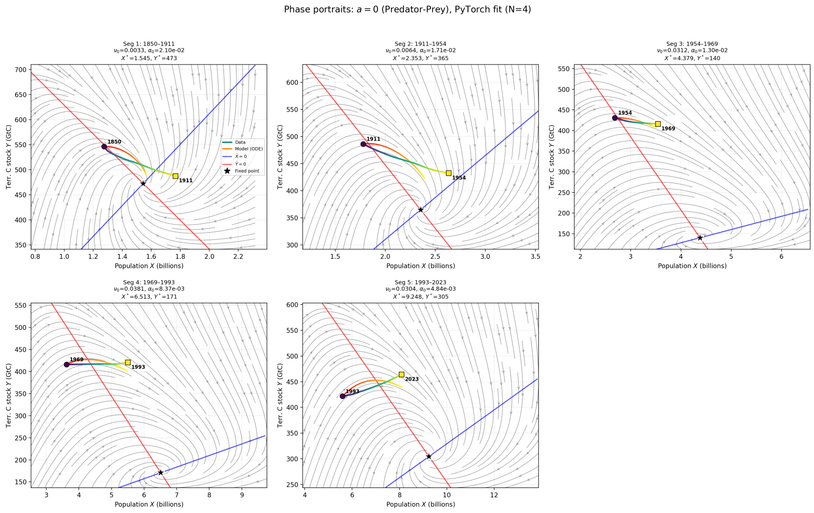


Figure 9: Phase portraits for the Predator–Prey model ($a = 0$) across five segments ($N_{\text{bp}} = 4$). Vector field (arrows), $X$- and $Y$-nullclines (dashed), non-trivial fixed point (star), observed data (circles), and model trajectory (solid line).

#### 5 Extended Model — Unified Three-Interaction Framework

##### 5.1 Model formulation

To assess whether empirical data favor a single interaction mechanism or a combination thereof, we formulate an extended model that simultaneously incorporates all three functional forms:

$$\frac{dX}{dt} = r_0 \, X \left(1 - \frac{X}{\nu_0 \, Y}\right) \tag{42}$$

$$\frac{dY}{dt} = \beta_0 \, Y \left(1 - \frac{Y}{K}\right) - \alpha_0 \, X \, Y - \alpha_1 \, X - \alpha_2 \, \frac{X}{Y} \tag{43}$$

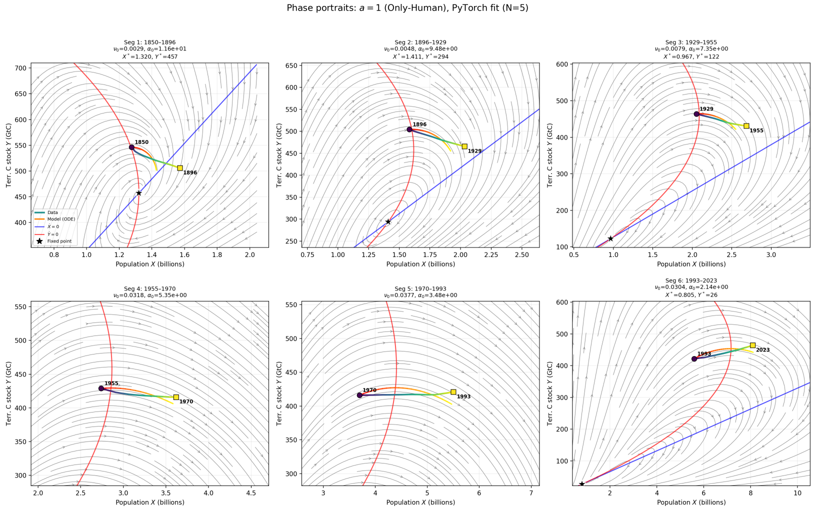


Figure 10: Phase portraits for the Only-Human model ($a = 1$) across six segments ($N_{\text{bp}} = 5$).

where $\alpha_0$ (Predator–Prey), $\alpha_1$ (Only-Human), and $\alpha_2$ (Supply–Demand) are simultaneously active. The piecewise-constant framework now has four parameters per segment $(\nu_0, \alpha_0, \alpha_1, \alpha_2)$, yielding

$$n_{\text{params}} = 5\,N_{\text{bp}} + 4. \tag{44}$$

###### 5.1.1 Fixed-point derivation

The $X$-nullcline gives $X^* = \nu_0\,Y^*$. Substituting into $dY/dt = 0$:

$$\beta_0\,Y^*\left(1 - \frac{Y^*}{K}\right) - \alpha_0\,\nu_0\,Y^{*2} - \alpha_1\,\nu_0\,Y^* - \alpha_2\,\nu_0 = 0. \tag{45}$$

Dividing by $Y^*$ (for $Y^* \neq 0$) and rearranging yields a quadratic in $Y^*$:

$$A\,Y^{*2} - B\,Y^* + C = 0, \tag{46}$$

where

$$A = \frac{\beta_0}{K} + \nu_0\,\alpha_0, \qquad B = \beta_0 - \nu_0\,\alpha_1, \qquad C = \nu_0\,\alpha_2. \tag{47}$$

The solutions are

$$Y^* = \frac{B \pm \sqrt{B^2 - 4AC}}{2A}, \qquad X^* = \nu_0\,Y^*. \tag{48}$$

Existence of real coexistence equilibria requires $B^2 - 4AC \geq 0$ and $Y^* > 0$.

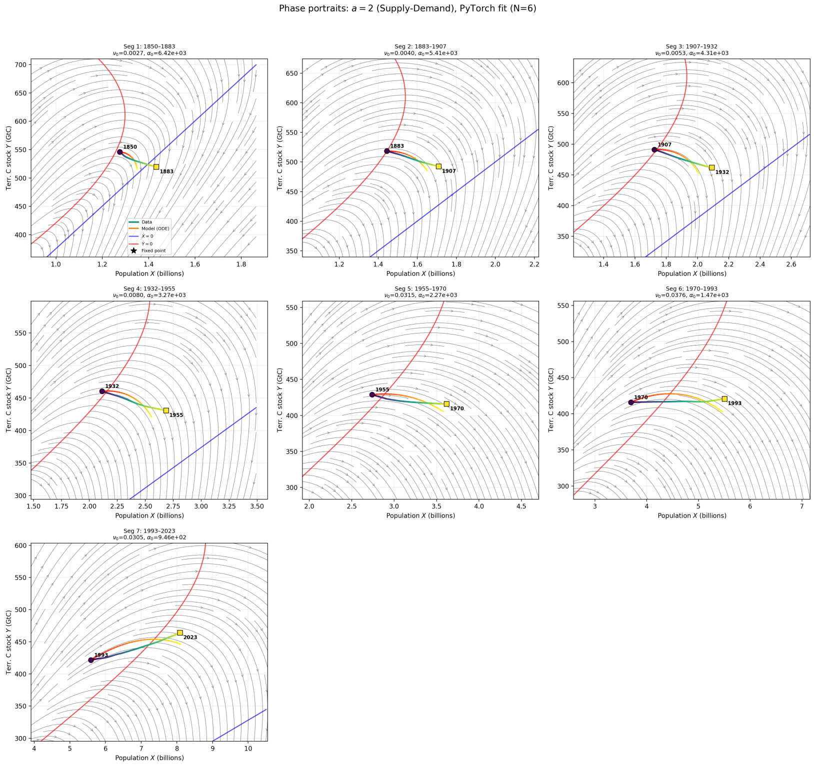


Figure 11: Phase portraits for the Supply–Demand model ($a = 2$) across seven segments ($N_{\mathrm{bp}} = 6$).

#### 5.2 Fitting results

##### 5.2.1 Multi-seed initialization

Because the extended model has a larger parameter space, we employ a multi-seed initialization strategy: three seeds derived from the best-fit individual models ($a = 0, 1, 2$), plus one "split" seed that distributes the total consumption from the best individual fit among the three $\alpha$ channels with weights $(0.5, 0.3, 0.2)$. The split seed is used by default as it consistently yields the best results.

##### 5.2.2 Best-fit summary

The extended model uses $N_{\text{bp}} = 6$ (the maximum Kneedle-detected value across the three individual models), yielding 7 segments. Table 4 compares the extended model against the three individual models.

Table 4: Comparison of all four models. The extended model uses $N_{\text{bp}} = 6$ (max Kneedle across individual cases) and achieves the lowest RSS.

| **Model** | $N_{\text{bp}}$ | $n_{\text{params}}$ | RSS |
|---|---|---|---|
| Predator–Prey ($a = 0$) | 4 | 14 | 0.0414 |
| Only-Human ($a = 1$) | 5 | 17 | 0.0313 |
| Supply–Demand ($a = 2$) | 6 | 20 | 0.0252 |
| **Extended** | 6 | 34 | **0.0177** |

The extended model achieves the lowest RSS (0.0177) across all four models, with breakpoints at 1893, 1920, 1941, 1956, 1971, and 1993.

#### 5.3 Phase portraits

#### 5.4 Consumption decomposition

To compare the relative importance of the three interaction mechanisms, we convert the dimensionally incompatible $\alpha$ parameters into effective consumption rates ($\text{GtC}\,\text{yr}^{-1}$) evaluated at the segment-mean state $(\bar{X}, \bar{Y})$:

$$C_0 = \alpha_0\,\bar{X}\,\bar{Y} \quad \text{(Predator–Prey)}, \tag{49}$$

$$C_1 = \alpha_1\,\bar{X} \quad \text{(Only-Human)}, \tag{50}$$

$$C_2 = \alpha_2\,\frac{\bar{X}}{\bar{Y}} \quad \text{(Supply–Demand)}. \tag{51}$$

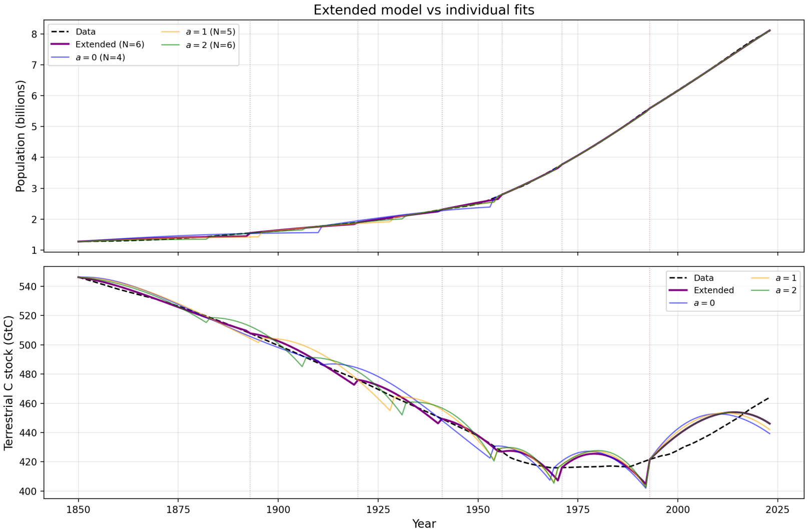


Figure 12: Best fit of the extended model ($N_{\mathrm{bp}} = 6$). Observed data (dashed) and model trajectories (solid) for $X(t)$ and $Y(t)$.

The fractional share of each mechanism in segment $j$ is $f_i^{(j)} = C_i^{(j)}/(C_0^{(j)} + C_1^{(j)} + C_2^{(j)})$ for $i = 0, 1, 2$.

Breakpoint attribution is performed analogously, evaluating $\Delta C_i$ at the breakpoint state $(X_{b_j}, Y_{b_j})$ to quantify which mechanism drives the change in total consumption across each transition.

### 6 Residual Bootstrap Analysis

#### 6.1 Methodology

To assess the uncertainty of parameter estimates and consumption shares in the extended model, we perform a residual bootstrap with $B = 300$ replicates. The procedure is as follows:

1. Compute the best-fit model trajectory via the multiple-shooting forward pass, obtaining model predictions $X_{\mathrm{mod}}(t)$ and $Y_{\mathrm{mod}}(t)$.

2. Compute residuals: $r_X(t) = X(t) - X_{\mathrm{mod}}(t)$ and $r_Y(t) = Y(t) - Y_{\mathrm{mod}}(t)$.

3. For each bootstrap replicate $b = 1, \ldots, B$:

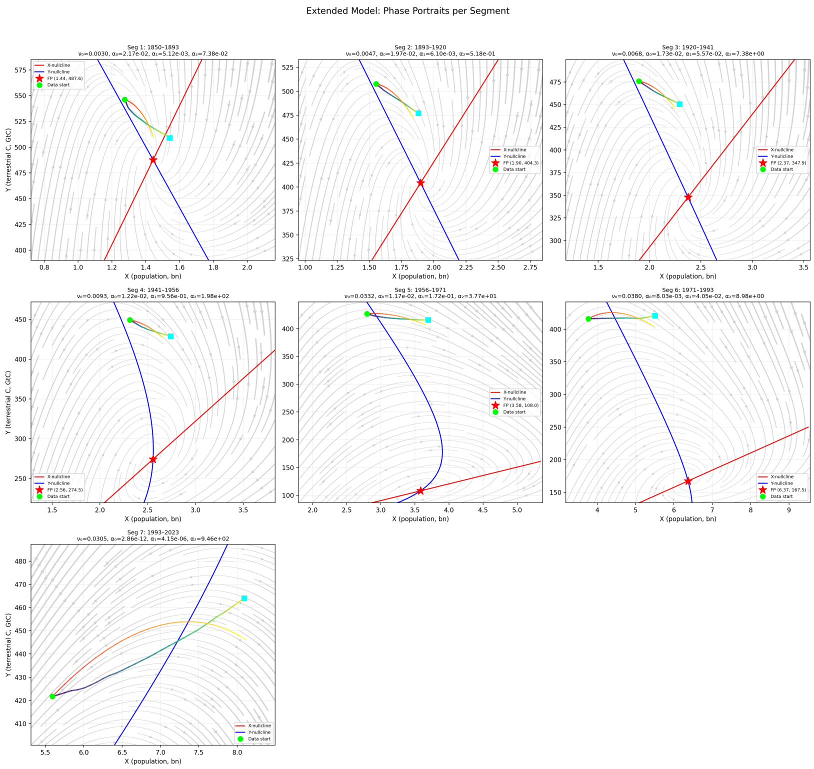


Figure 13: Phase portraits for the extended model across seven segments ($N_{\text{bp}} = 6$), showing vector field, nullclines, coexistence fixed points, observed data, and model trajectories.

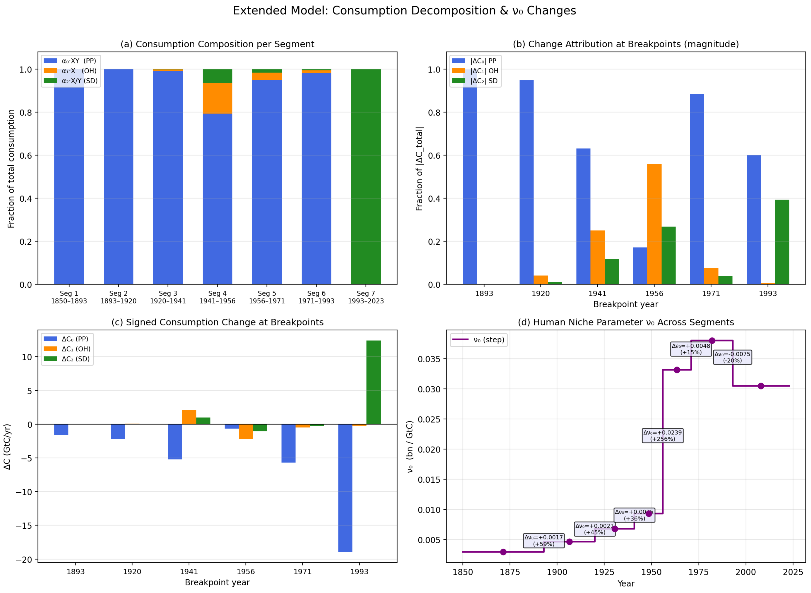


Figure 14: Consumption decomposition for the extended model: effective consumption rate ($\mathrm{GtC\,yr^{-1}}$) attributed to each mechanism (PP, OH, SD) per segment, evaluated at segment-mean state.

(a) Resample residual indices with replacement to construct bootstrap residuals $r_X^{*(b)}(t)$ and $r_Y^{*(b)}(t)$.

(b) Generate bootstrap data: $X^{*(b)}(t) = X_{\text{mod}}(t)+r_X^{*(b)}(t)$ and $Y^{*(b)}(t) = Y_{\text{mod}}(t) + r_Y^{*(b)}(t)$.

(c) Re-fit the extended model with $N_{\text{bp}} = 6$ and fixed breakpoints using the split-seed initialization.

4. Collect parameter estimates and derived quantities (consumption shares) across all replicates.

Breakpoint locations are held fixed at the best-fit values to focus uncertainty quantification on the interaction parameters. Parallel execution is achieved via Python `multiprocessing`.

#### 6.2 Results

All 300 replicates converged successfully. Confidence intervals (2.5th–97.5th percentile) for parameters and consumption shares are reported in Figure 15.

#### 6.3 Dominance frequency analysis

For each segment, we compute the percentage of bootstrap replicates in which each mechanism has the largest consumption share. Table 5 reports these frequencies together with the fraction of replicates where each mechanism exceeds 50% of total consumption—a stronger test than mere dominance.

Table 5: Dominance frequency (%): fraction of $B = 300$ bootstrap replicates in which each mechanism has the largest consumption share per segment, and fraction where each mechanism's share exceeds 50% of total consumption.

| | **Seg 1** 1850–93 | **Seg 2** 1893–1920 | **Seg 3** 1920–41 | **Seg 4** 1941–56 | **Seg 5** 1956–71 | **Seg 6** 1971–93 | **Seg 7** 1993–2023 |
|---|---|---|---|---|---|---|---|
| PP dominant | 86.7 | 85.0 | 82.3 | 84.3 | 93.7 | 95.7 | 16.7 |
| OH dominant | 2.3 | 3.0 | 3.3 | 6.7 | 1.7 | 2.7 | 1.7 |
| SD dominant | 11.0 | 12.0 | 14.3 | 9.0 | 4.7 | 1.7 | 81.7 |
| PP share $> 50\%$ | 84.7 | 78.0 | 74.0 | 56.7 | 63.7 | 76.3 | 15.3 |
| OH share $> 50\%$ | 0.0 | 0.0 | 0.0 | 0.0 | 0.0 | 0.0 | 0.0 |
| SD share $> 50\%$ | 6.3 | 6.0 | 7.3 | 1.3 | 1.0 | 0.0 | 77.7 |

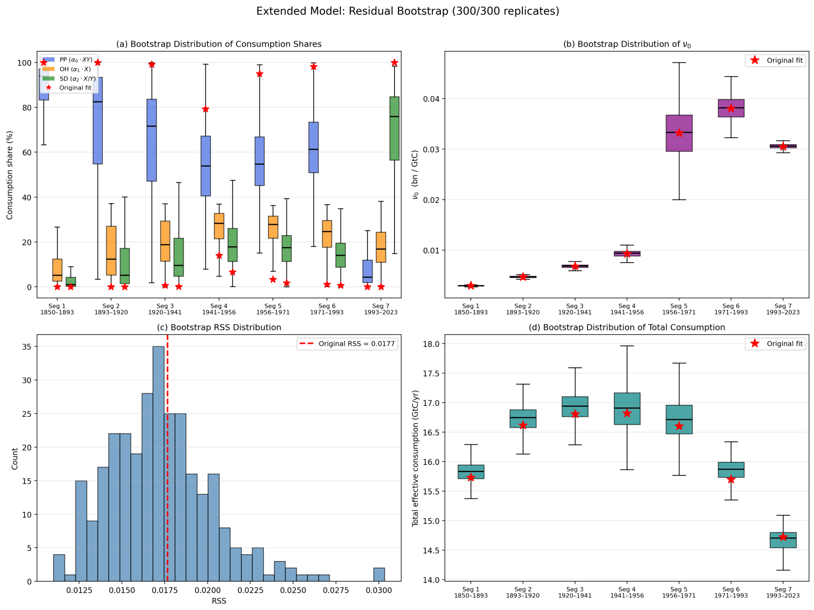


Figure 15: Bootstrap analysis of the extended model ($B = 300$ replicates). Parameter estimates and consumption share confidence intervals (2.5th–97.5th percentile) per segment.

The Predator–Prey (PP) interaction robustly dominates in segments 1–6 (1850–1993), appearing as the largest consumption mechanism in 82–96% of bootstrap replicates. The PP share exceeds half of total consumption in 57–85% of cases across these segments. Segment 7 (1993–2023) is markedly different: the Supply–Demand (SD) mechanism appears dominant in 82% of replicates, with SD share exceeding 50% in 78% of replicates.

The strongest result of the analysis concerns the Only-Human (OH) mechanism: it never accounts for a majority of consumption in any segment across any of the 300 bootstrap replicates (OH share $> 50\% = 0\%$ in all segments). This rules out constant per-capita consumption ($\alpha_1 X$) as a standalone dominant mechanism with full statistical confidence.

These results should be interpreted with the following caveats. The three $\alpha$ parameters are structurally non-identifiable in the extended model when evaluated at a single state point, and the confidence intervals on individual $\alpha$ estimates are notably wider in segment 7 than in earlier segments (cf. Figure 15). The apparent shift to SD dominance in the most recent period may therefore partly reflect estimation uncertainty. Additionally, the bootstrap captures statistical but not structural model uncertainty.

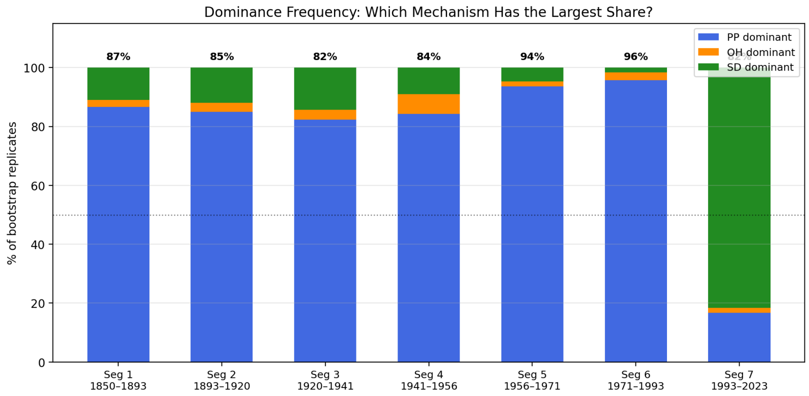


Figure 16: Dominance frequency across bootstrap replicates: fraction of $B = 300$ replicates in which each mechanism (PP, OH, SD) has the largest consumption share, per segment.
